# Cingulate-centered flexible control: physiologic correlates and enhancement by internal capsule stimulation

**DOI:** 10.1101/2025.10.15.682151

**Authors:** Jaejoong Kim, Alik S. Widge

## Abstract

Flexible cognitive control requires rapid adjustment when current demands diverge from the brain’s predictions. Deficits in control adjustment are prominent in psychiatric disorders, yet the neural mechanisms supporting such adjustment and their modifiability remain unclear. We analyzed two intracranial electroencephalography datasets, one with brief internal capsule stimulation (ICS), to identify a human circuit mechanism for flexible adjustment after control-demand change. We further used two clinical internal capsule deep brain stimulation (IC DBS) cohorts to validate our proposed mechanism. Across intracranial datasets, phase-amplitude coupling (PAC) anchored to the theta phase of right rostral anterior cingulate cortex (rACC-R), particularly theta–gamma coupling with dorsolateral prefrontal cortex and dorsal ACC, was associated with faster performance specifically when control demand changed. This effect generalized across datasets and tasks and was robust across analytic pipelines. An adaptive drift-diffusion model accounted for this adjustment cost in terms of unsigned control prediction error. In Dataset 1, brief ICS selectively enhanced the same rACC-R theta-centered PAC configuration on control-state-change trials, and computational modeling indicated that stimulation improved control flexibility under high-mismatch conditions. In a primarily treatment-resistant depression IC DBS cohort, stimulation-induced enhancement of the flexibility-related behavioral parameter, rather than a more general-control parameter, was associated with clinical response (N = 14; AUC = 0.90). In a separate treatment-resistant obsessive-compulsive disorder IC DBS cohort, clinically selected stimulation settings preferentially increased the same flexibility-related parameter across all patients (N = 5). These findings identify rACC-centered theta-phase coordination as a candidate human mechanism for flexible adjustment, show that capsule stimulation can augment this mechanism when flexibility is required, and suggest a causal link between flexibility-related behavioral and neural signals and psychiatric symptoms.

## Introduction

The flexible deployment of cognitive control is essential for adaptive functioning in dynamic environments where cognitive resources are limited ^1–4^. When task demands change unexpectedly, the brain must rapidly suppress now-irrelevant control settings and reconfigure toward a more appropriate state ^4^. For example, imagine settling into a café anticipating a quiet workspace for a difficult task. When unexpected loud conversations erupt nearby, your brain must rapidly redirect resources toward suppressing these distractions. Once the noise subsides, cognitive control allows you to flexibly shift those resources back to your primary task, optimizing productivity. Impairments in this process (cognitive rigidity) are prominent in psychiatric disorders such as treatment-resistant depression (TRD) and obsessive–compulsive disorder (OCD), where patients struggle to shift away from maladaptive thought and behavioral patterns ^5,6^. Enhancing control flexibility could therefore alleviate clinical symptoms, yet conventional treatments including medications and psychotherapies have not been shown to improve it specifically. This shortfall stems partly from our limited understanding of the neural mechanisms underpinning control flexibility and from the lack of efficient ways to modulate those mechanisms.

A central player in many models of cognitive control is the anterior cingulate cortex (ACC), which may monitor ongoing performance and signal when adjustments are required ^7^. Various accounts attribute different roles to the ACC—detecting conflict ^8^, predicting task difficulty ^7^, or encoding an error signal between predicted and actual demands ^3,9^. Across these frameworks, however, a common idea is that the ACC detects when the current control state no longer matches task demands and helps trigger the flexible adjustment of control ^9–13^. Several computational accounts describe this ACC-related signal in terms of valence-neutral mismatch or surprise ^12,13^ which has been formalized as an unsigned control prediction error (CPE). What remains unclear, however, is how the human ACC implements this transition from mismatch detection to efficient control reconfiguration at the network level.

One candidate mechanism is phase-amplitude coupling (PAC) within distributed control networks ^14^. Computational and animal work suggests that ACC theta oscillations (4-8 Hz) may organize flexible control by modulating higher-frequency oscillations (HFO) in downstream regions, particularly lateral prefrontal cortex (LPFC), during periods when reconfiguration is required ^15–17^. More broadly, human work on midfrontal theta has proposed that medial prefrontal theta activity may coordinate adaptive control by facilitating communication across distributed brain regions ^18^. Consistent with this idea, ACC theta-phase coupling (5-10 Hz) with gamma-frequency power (35-55 Hz) in LPFC has been associated with attentional shifting in macaques ^17^, and related theoretical work implicates orbitofrontal and hippocampal interactions in model-based control updating ^19,20^. There remains limited direct human evidence that PAC between ACC theta phase and HFO of other control regions, including LPFC, supports efficient adjustment after control-demand change. Clarifying this mechanism would help bridge computational theories of adaptive control with human network physiology and could identify neural signals that are modifiable for therapeutic purposes.

One potential way to modulate ACC-mediated control flexibility is deep brain stimulation (DBS). DBS targeting fronto-subcortical circuits has been used to relieve symptoms in TRD and OCD ^21–23^. Although the precise mechanism is still unclear, DBS can modulate neural synchrony across distributed brain networks ^24,25^. For example, stimulation in the subthalamic nucleus can suppress pathological beta–gamma PAC across connected regions ^24^, while stimulation of subgenual cingulate cortex may enhance beneficial interactions such as theta–gamma PAC linked to cognitive processing ^25^. Particularly, DBS of the internal capsule (IC), a dense white-matter tract with fibers originating in ACC, LPFC, and OFC, can engage multiple nodes of the cognitive control network simultaneously ^1,26,27^. Consistent with this broad network engagement, we recently showed that IC DBS improves response speed on cognitive control tasks in both healthy individuals and patients with TRD/OCD, without compromising accuracy ^28,29^. Moreover, computational modeling indicates that this enhancement stems from a faster rate of evidence accumulation (conflict resolution), in rodents and humans ^30^. However, it remains unclear whether stimulation specifically enhances the flexible adjustment component of control, or instead improves a more general aspect of cognitive control that is not specifically related to flexibility. This distinction is important because the same computational framework can be used to test whether stimulation preferentially enhances adjustment when control-demand mismatch is high, rather than merely increasing context-independent control. It is likewise unclear whether stimulation-induced gains in cognitive control and/or its flexible deployment are related to IC DBS’ clinical effects.

Here, we test three hypotheses across multiple datasets. First, using two intracranial EEG datasets, we test whether flexible adjustment after control-demand change in cognitive control tasks depends on ACC-centered PAC, particularly theta phase in ACC coupled to HFO in putative downstream control regions identified in prior literature. Second, using internal capsule stimulation in Dataset 1, we test whether this flexibility mechanism can be causally enhanced specifically when control-demand mismatch is high. Third, using two clinical IC DBS cohorts, we test whether the flexibility-related component of stimulation is clinically relevant: in Dataset 3, by asking whether DBS-induced flexibility enhancement is associated with therapeutic response, and in Dataset 4, by asking whether clinically effective IC DBS settings in TR-OCD preferentially enhance the same flexibility-related component.

### Datasets and Tasks

We studied control flexibility across four complementary datasets. Dataset 1 involved epilepsy patients performing the Multi Source Interference Task (MSIT) while undergoing iEEG recording with and without brief ICS, enabling us to identify a network-level mechanism underlying flexibility. Dataset 2 replicated and extended these findings across three cognitive tasks (Table 1) in an independent cohort collected outside our laboratory. Dataset 3 tested whether DBS-induced improvements in control flexibility predicted clinical outcomes in patients with treatment-resistant depression or OCD. Dataset 4 tested whether clinically selected IC DBS settings in a separate TR-OCD cohort preferentially enhanced the same flexibility-related computational component. Full task and sample details are provided in the Methods.

**Table 1.**
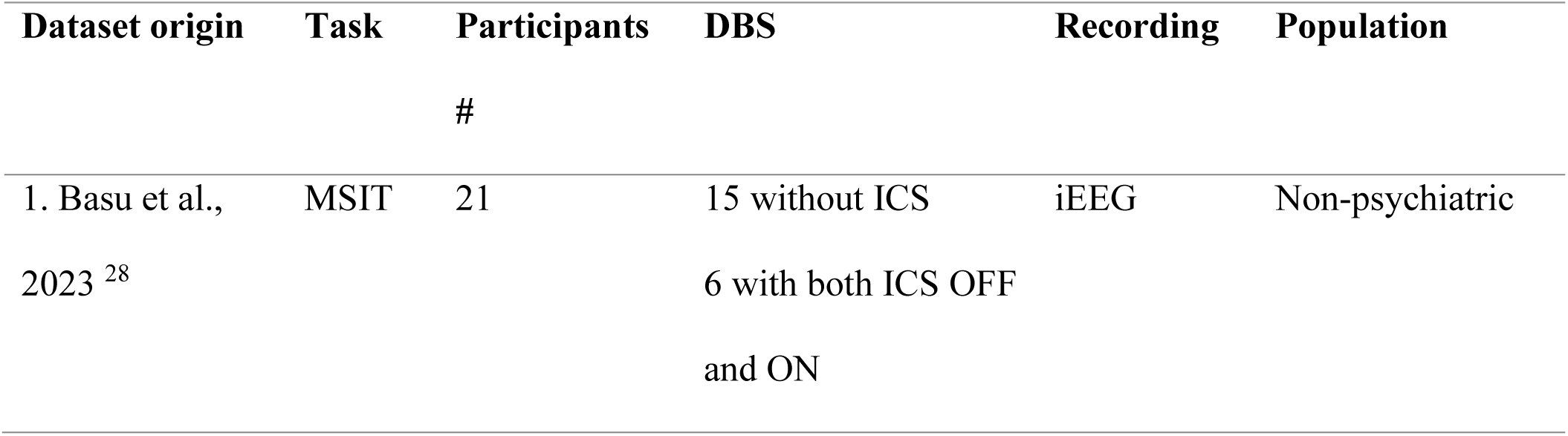
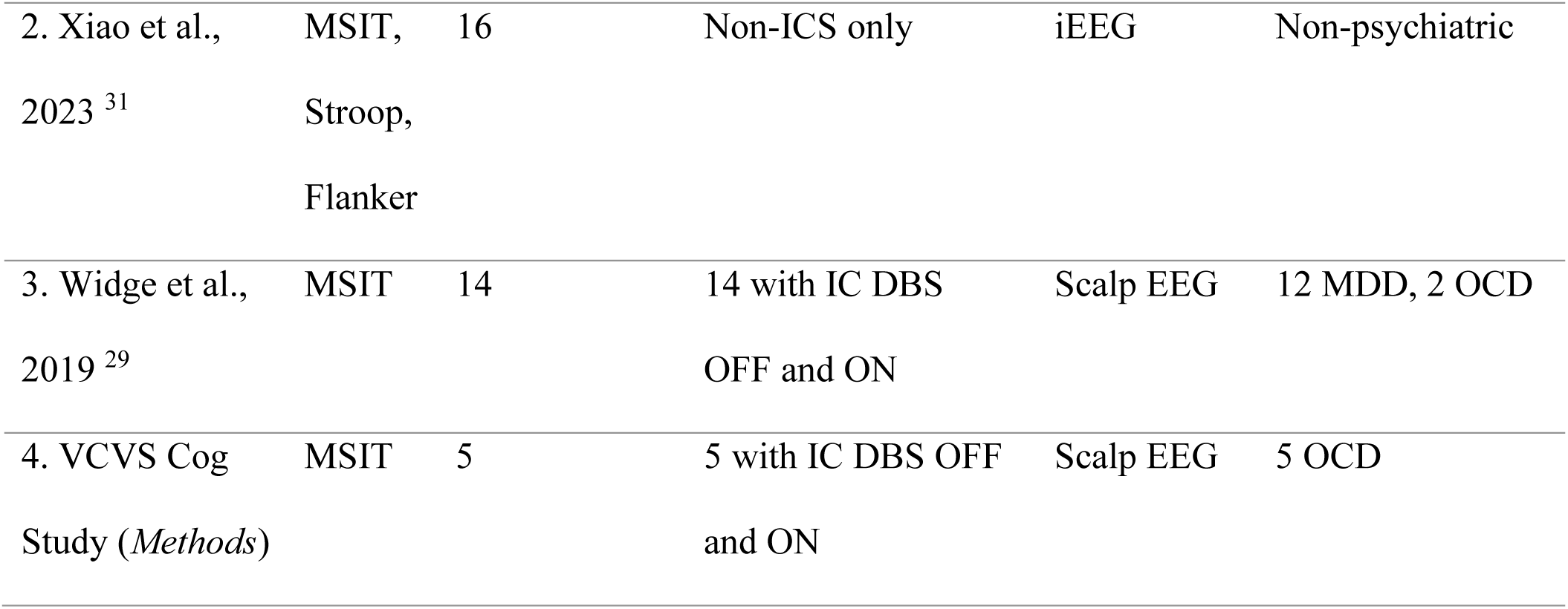

In all four datasets, participants performed MSIT or analogous conflict-based tasks (Stroop, Flanker in Dataset 2). MSIT requires participants to identify the unique number in a three-digit display (e.g., “2 2 1”) by pressing a button matching identity, not its spatial position (Figure 1A). Conflict trials introduce interference by mismatching the correct number’s identity and location (e.g., “1” appearing in the third position), while Non-conflict trials have no such mismatch. Across all tasks, trials are presented in a pseudorandom sequence such that Conflict and Non-conflict trials alternate unpredictably.

**Figure 1.**
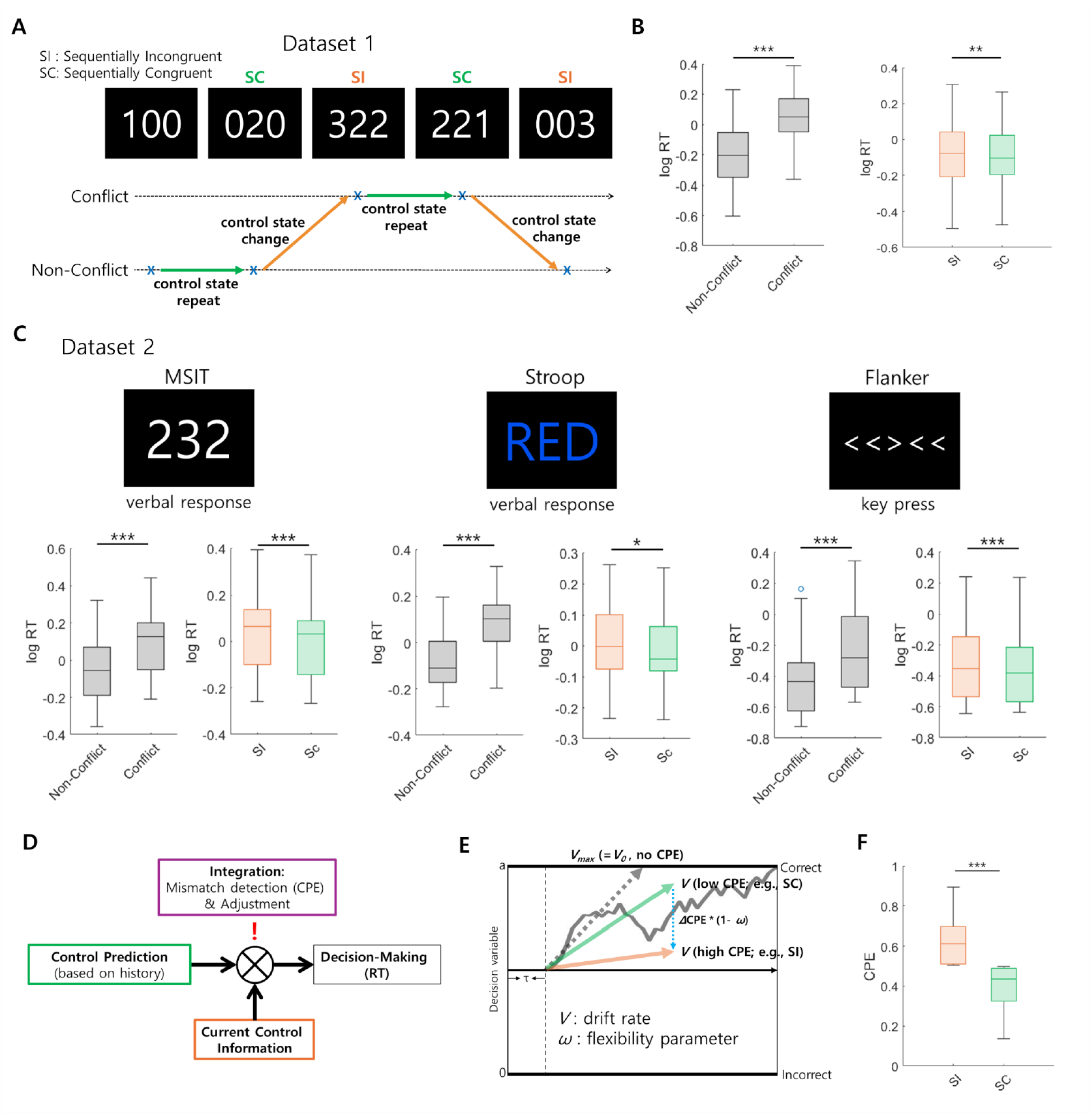
Behavioral Evidence for Control Prediction Errors (CPEs) and Their Computational Modeling. (A) Schematic of the Multi-Source Interference Task (MSIT) and the control-state change during the task. Sequentially incongruent (SI) trials denote a change in control state relative to the previous trial, whereas sequentially congruent (SC) trials denote repetition of the prior control state. (B) In Dataset 1 (N=21), conflict trials elicited longer reaction times (RTs) than non-conflict trials, and SI trials elicited significantly longer RTs than SC trials, indicating an adjustment cost when control demand changed. (C) This SI>SC RT effect, as well as Conflict>Non-conflict RT effect, was replicated in Dataset 2 across MSIT, Stroop, and Flanker tasks. (D) Schematic of the adaptive drift-diffusion model (adaptive DDM), in which predicted control demand is updated from recent trial history and compared with current control demand; the resulting mismatch is formalized as control prediction error (CPE). (E) In the adaptive DDM, higher mismatch reduces drift rate, whereas the flexibility parameter, ω, attenuates this mismatch-related cost. (F) Model-derived unsigned CPE was higher on SI than SC trials, showing that the task-level SI/SC contrast corresponds to greater control-demand mismatch in the computational model. Boxplots show median and interquartile range; whiskers denote the data range. * p<0.05, ** p<0.01, *** p<0.001.

This task structure produces trial-to-trial changes in control demand. Our primary interest is in how the brain adjusts when the current trial requires a different control state from the one just engaged on the previous trial. At the task level, we operationalize this as sequential incongruency (SI) versus sequential congruency (SC): SI trials are those in which the current trial type differs from the previous trial, whereas SC trials repeat the previous control state. We focus on this SI–SC contrast because it captures whether control demand changed or repeated from one trial to the next, which is the task-level feature most relevant to our central question about the neural mechanism of rapid adjustment after control-state change. Although this framing is related to, but not identical with, the classical Gratton effect ^32–34^, which focuses on the interaction between previous-trial conflict and current-trial conflict, our focus here is the neural signal that supports efficient adjustment when the control state changes; we discuss this relationship further in the Supplementary Information (see *Supplementary Results*).

## Results

### Behavioral evidence for adjustment after control-demand change

In Dataset 1 (Table 1), conflict trials had longer RTs than non-conflict trials (all *p* < 0.001; Figure 1B). More importantly for the present study, participants also showed longer log RTs on SI trials than on SC trials (*t* = 2.97, *p* = 0.002, in linear mixed-effects regression, LMER; Figure 1B), indicating an additional adjustment cost when control demand changed from the previous trial. In Dataset 2, this finding replicated across the MSIT, Stroop, and Flanker tasks (all *p* < 0.05; Figure 1C).

We next asked whether this RT slowing after control-demand change could be explained computationally (see *Methods*). Following prior work that frames flexible cognitive control in terms of trial-by-trial prediction of changing control demands ^3,8,35^, we tested an adaptive drift-diffusion model (adaptive DDM) that extends the basic DDM by incorporating an internal estimate of conflict probability updated across trials. In this model, mismatch between predicted and actual control demand modulates the drift rate, such that larger mismatch transiently slows evidence accumulation and therefore lengthens RT. The flexibility parameter, ω, captures how strongly this mismatch affects drift rate: larger ω values reduce the behavioral impact of mismatch and thus operationalize greater control flexibility (Supplementary Methods). Fitting this adaptive DDM to trial-by-trial RT distributions and accuracy significantly outperformed a basic DDM in Dataset 1 (ΔDIC = –2502; Supplementary Figure 1). This result was replicated in the MSIT in Dataset 2 (ΔDIC = –127) and also held in the Stroop and Flanker tasks (ΔDIC = –33 and –146, respectively; Supplementary Figure 1), indicating that a model incorporating dynamic control-demand mismatch explained behavior better than a fixed-drift-rate model across datasets.

Consistent with this account, simulated output from the adaptive DDM showed greater RT prolongation under high-mismatch conditions in an agent with low ω (*p* < 0.001, 10,000 simulations; Supplementary Figure 1), whereas an agent with high ω was substantially less affected (*p* = 0.407; Supplementary Figure 1). Model-derived control prediction error (CPE) was also higher on SI than SC trials (average CPE: 0.63 in SI trials vs 0.39 in SC trials; *p* < 0.001; Fig. 1H), indicating that the task-level contrast of control-demand change versus repetition was associated with larger unsigned mismatch in the computational model. We focus on unsigned CPE because our primary question concerns the magnitude of control-state mismatch requiring reconfiguration, rather than whether control demand increased or decreased (see Supplementary materials). Thus, the adaptive DDM provides a mechanistic account of why RT slows after control-demand change and motivates the later use of SI versus SC as a task-level contrast for studying the neural and stimulation-related mechanisms of flexible adjustment.

### Neural mechanisms supporting flexible adjustment after control-demand change (Dataset 1, ICS OFF)

Having established that reaction time slows when control demand changes from one trial to the next, we next examined the neural mechanisms that support efficient adjustment to such changes in Dataset 1 (stimulation OFF condition only; Table 1). These 21 patients had intracranial EEG coverage in key frontal, cingulate, and hippocampal regions implicated in flexible cognitive control, including ACC (rostral and dorsal subregions), dlPFC, lateral orbitofrontal cortex (lOFC), and hippocampus (Figure 2A) ^7,19^. Our primary question was which neural signals support efficient adjustment when control demand changes from one trial to the next. Using SI versus SC as the task-level contrast, we asked which local or network-level signals were specifically associated with faster performance when the control state had to be reconfigured, rather than with general performance across both trial types.

**Figure 2.**
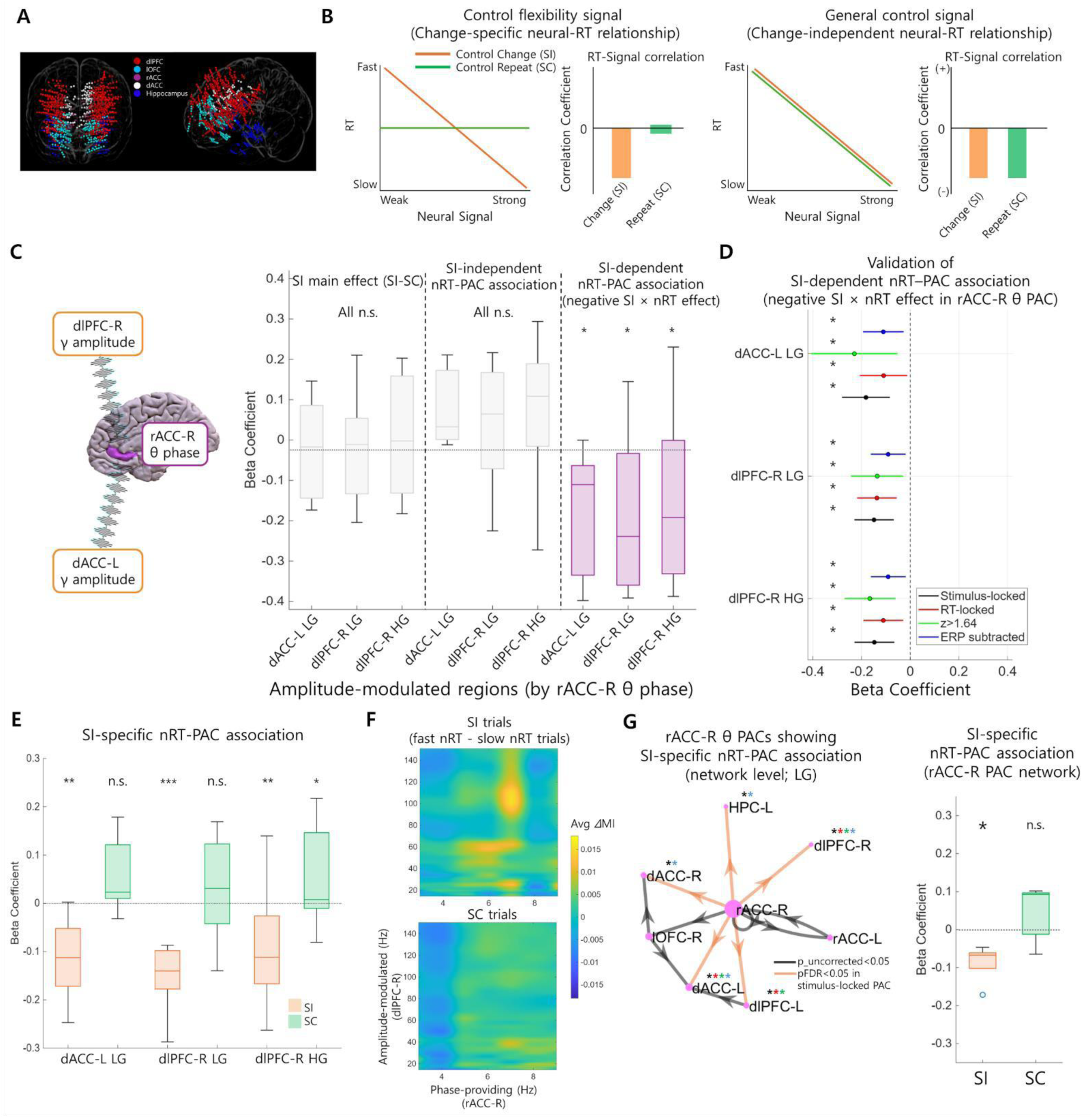
Neural mechanisms supporting flexible control (Dataset 1; intracranial EEG, DBS OFF). (A) Anatomical electrode coverage for the 21 epilepsy patients in Dataset 1. (B) Conceptual schematic of the neural test. A control flexibility signal is expected to show a stronger negative neural-RT relationship when control demand changes (SI) than when it repeats (SC), whereas a more general control signal should show a similar neural-RT relationship across SI and SC trials. (C) LMER results for the three most robust rACC-R theta-centered PAC pairs—rACC-R theta–dACC-L LG, rACC-R theta–dlPFC-R LG/HG. These PACs showed no SI main effect and no SI-independent PAC–nRT association, but did show a significant SI-dependent PAC–nRT association (negative SI × nRT effect). nRT, normalized log RT, was computed by z-scoring log RT within conflict and non-conflict trials. (D) Validation of the same three PAC effects across complementary PAC pipelines: stimulus-locked PAC, RT-locked PAC, PAC restricted to electrode pairs with significant trial-average PAC (z > 1.64), and condition-specific ERP-subtracted PAC. (E) Post-hoc within-condition analyses showing that the same three PACs were negatively associated with nRT on SI trials but not on SC trials. (F) Representative rACC-R-centered comodulograms illustrating stronger theta– gamma coupling on fast-versus slow-response SI trials relative to SC trials. (G) Network-level summary of rACC-R theta-centered PAC, derived by principal component analysis (PC1), showing that stronger rACC-R PAC network coupling was associated with faster nRT specifically on SI trials. Boxplots show median and interquartile range; whiskers denote the data range. Boxplots show median and interquartile range; whiskers denote the data range. In forest plots, circles indicate beta coefficients and horizontal lines indicate 95% confidence intervals. Asterisks indicate FDR-corrected significance unless otherwise noted. * p<0.05, ** p<0.01, *** p<0.001.

We therefore asked whether the relationship between trial-by-trial neural activity and normalized log RT (nRT; see Methods) differed between trials in which control demand changed versus repeated (Figure 2B). We used normalized log RT (nRT), computed by z-scoring log RT within conflict and non-conflict trials, to remove large conflict-driven RT differences and test whether neural activity tracked relatively faster or slower responses within each control-demand state. A neural signal supporting flexible adjustment should be associated with faster responses specifically when control demand changes, because this is when prior control settings must be reconfigured ^36–38^ and resources must be redirected toward the currently appropriate control state ^4^ (Figure 2B, left). In contrast, a more general control signal should facilitate performance across both changed and repeated control states, without a selective interaction with SI versus SC (Figure 2B, right). Such a signal may reflect sustained attention or an overall upregulation of control that improves response speed regardless of change ^8,39^. Thus, our key test was whether a neural signal showed a stronger negative relationship with nRT on SI than on SC trials, yielding a negative SI × nRT interaction (Figure 2B, left).

### rACC-R theta–gamma PAC with dlPFC/dACC: a neural signature of flexible adjustment

To investigate whether phase-amplitude coupling (PAC) underlies flexible adjustment, we computed trial-by-trial PAC between theta oscillations and the amplitude envelope of high-frequency oscillation (HFO; including beta (15-30 Hz), low-*γ* (30-50 Hz; LG), and high-*γ* (70-150 Hz; HG)) for all electrode pairs during the 0.1–1.4 s post-stimulus window. We then used linear mixed-effects regression (LMER; see Methods) to test whether the relationship between trial-by-trial PAC for each regional pair (10 × 10 pairs) and nRT showed the predicted negative SI × nRT interaction.

Strikingly, significant negative SI × nRT interactions emerged in multiple target regions only when right rostral ACC (rACC-R) served as the theta-phase source (pFDR < 0.05, corrected for 10 × 10 regional PAC pairs; Figure 2C; Supplementary Figure 2). No other phase source showed this pattern, and there was no condition-independent PAC–nRT correlation (Figure 2C; Supplementary Figure 2). Thus, rACC-R-centered PAC showed the predicted negative SI × nRT interaction, indicating that the PAC–behavior relationship differed between changed and repeated control states in a manner consistent with a flexible-adjustment signal rather than a more general performance-related signal.

To exclude the possibility that local rACC-R theta power itself was driving these effects, we repeated the nRT analyses using rACC-R theta power and found no SI × nRT interaction (pFDR > 0.05; Supplementary Figure 3). Instead, rACC-R theta power was more consistent with a general control signal, showing a condition-independent negative relationship with nRT (Supplementary Figure 3).

To further verify the robustness of the rACC-R PAC effects, we recomputed PAC using three complementary pipelines. First, we recomputed PAC time-locked to the response (RT-locked) rather than to stimulus onset (stimulus-locked). Second, we reran the model after restricting it to electrode pairs whose trial-average PAC exceeded z = 1.64 (“significant” trial-average PAC; criterion from Daume et al., 2024 ^40^; 500 trial-shuffled surrogates; see Methods). This approach risks missing electrode pairs that are weak on average yet become strongly coupled in specific contexts, such as fast-response SI trials. Finally, to exclude the possibility that the observed PAC effects were driven by ERP differences between SI and SC conditions, we recomputed PAC after removing the average ERP for each condition (condition-specific ERP-subtracted PAC). Theta–gamma PAC between rACC-R and dlPFC-R (theta–LG and theta–HG), as well as theta–LG PAC between rACC-R and dACC-L which are two hubs of the conventional cognitive control network ^41,42^, survived the most conservative criterion by showing the same negative interaction pattern across all four methods (all pFDR < 0.05, corrected for the 18 region–frequency pairs showing significant negative interactions in the original stimulus-locked analysis; Figure 2D; Supplementary Figure 2). The rACC-R–dlPFC-L theta–gamma coupling survived in three of the four methods (Supplementary Figure 2).

Notably, the three PACs that survived the most conservative criterion across all four pipelines — rACC-R theta–dlPFC-R LG PAC, rACC-R theta–dlPFC-R HG PAC, and rACC-R theta–dACC-L LG PAC — were not specific to one transition direction. When SI trials were decomposed into conflict → non-conflict and non-conflict → conflict transitions (signed errors), the same negative interaction effects were observed in both directions (*Supplementary Results;* Supplementary Figure 4A). These results indicate that the rACC-centered PAC effect was not specific to either increasing or decreasing control demand, but instead reflected a broader process of reconfiguring control state when recent demand changed (unsigned error).

We next asked whether this task-level SI effect generalized to the model-derived mismatch signal (unsigned CPE). Using trial-by-trial unsigned CPE estimates from the adaptive DDM, we found a similar negative interaction pattern in the same three PACs — rACC-R theta–dlPFC-R LG PAC, rACC-R theta–dlPFC-R HG PAC, and rACC-R theta–dACC-L LG PAC — indicating that PAC–nRT coupling became more negative when unsigned CPE was high (*Supplementary Results;* Supplementary Figure 4C). We also tested whether these rACC-R PAC effects were better explained by signed, direction-specific mismatch than by unsigned mismatch. In both the SI-based and CPE-based analyses, the critical interaction remained associated with unsigned rather than signed mismatch (*Supplementary Results;* Supplementary Figure 4B-C). Together, these findings indicate that the relationship between rACC-R theta-centered PAC and behavior is better captured by non-directional control-demand change or mismatch than by a direction-specific signal. Computationally, this pattern is consistent with a mechanism in which the PAC–behavior relationship scales with the magnitude of detected control-demand mismatch—formalized here as unsigned CPE—rather than with the sign of that mismatch.

To clarify the form of the significant negative SI × nRT interaction observed in the original stimulus-locked PAC analysis, we next performed post-hoc analyses separately within SI and SC trials (see Methods). These analyses were intended to describe that interaction pattern, rather than to serve as independent discovery analyses. All three rACC-R theta–gamma PACs that showed the robust negative interaction pattern above — rACC-R–dACC-L theta–LG PAC and rACC-R–dlPFC-R theta–LG/HG PAC — were negatively correlated with nRT in SI trials (all pFDR < 0.05 in LMER, corrected across the 18 selected region– frequency pairs; Figure 2E), but not in SC trials (pFDR > 0.05 for dACC-L LG and dlPFC-R LG, with a positive correlation for dlPFC-R HG; Figure 2E–F). Thus, the significant SI × nRT interaction reflected stronger PAC–behavior coupling specifically when control demand changed.

Because similar negative SI × nRT interactions were observed across multiple PACs based on rACC-R theta phase (Supplementary Figure 2), we summarized the overall rACC-R PAC pattern using principal component analysis (PCA) to derive the dominant component reflecting rACC-R-theta-phase-centered network coordination (PC1, Supplementary Figure 5A; hereafter, the “rACC-R PAC network”; Figure 2G; see *Methods*). This network-level measure showed the same flexibility signature: stronger network coupling was associated with faster nRT in SI trials (t = –2.53, coefficient = –0.10, pFDR < 0.05 in LMER, corrected for four methods; Figure 2G), but not in SC trials (pFDR > 0.05; Figure 2G). This pattern was robust across the three validation methods above (Supplementary Figure 5B), underscoring the reliability of rACC-R-theta-centered coordination as a network correlate of flexible adjustment.

In addition to PAC analyses, we also examined local, non-phase-locked HFO power (*Supplementary Results;* Supplementary Figure 6). In contrast to the rACC-R PAC pattern, local HFO power, including bilateral dlPFC beta power, was more consistent with a general control signal, showing an nRT–power relationship without a selective SI interaction (Supplementary Figure 6). We also identified local HFO power encoding current conflict and conflict history (Supplementary Results; Supplementary Figure 6).

Together, these findings identify rACC-R theta-phase-based coordination, particularly theta–gamma PAC with dlPFC and dACC, as a neural signature of efficient adjustment when control demand changes from one trial to the next. This pattern was present across both directions of control-state change, generalized from the task-level SI contrast to model-derived unsigned mismatch, and was not better explained by signed or direction-specific formulations. These results support the view that rACC-R-centered PAC is associated with rapid reconfiguration of control state, rather than reflecting a more general performance-related signal.

### Replication and generalization of the rACC-R PAC network

Having established that rACC-R-centered PAC is associated with flexible adjustment in Dataset 1, we next asked whether this effect replicated and generalized beyond the MSIT and to an independently collected dataset. Dataset 2 ^31^ contained intracranial EEG from four participants with rACC-R electrodes who performed the MSIT, color-word Stroop, and Eriksen Flanker tasks. Using the same pipeline, we derived an rACC-R PAC network for each task (PC1 of rACC-R–theta-phase-based PAC), comprising coupling between rACC-R and right dlPFC as well as coupling within rACC-R itself. Across all three tasks, stronger rACC-R PAC network coupling was associated with faster nRT specifically on SI trials, but not on SC trials (all pFDR < 0.05 in LMER; Figure 3). Thus, the SI-specific negative PAC–nRT relationship replicated across datasets and generalized across multiple cognitive control paradigms. Consistent with the computational results above, this pattern supports the view that rACC-R-centered PAC tracks flexible adjustment when control demand changes from the previous trial.

**Figure 3.**
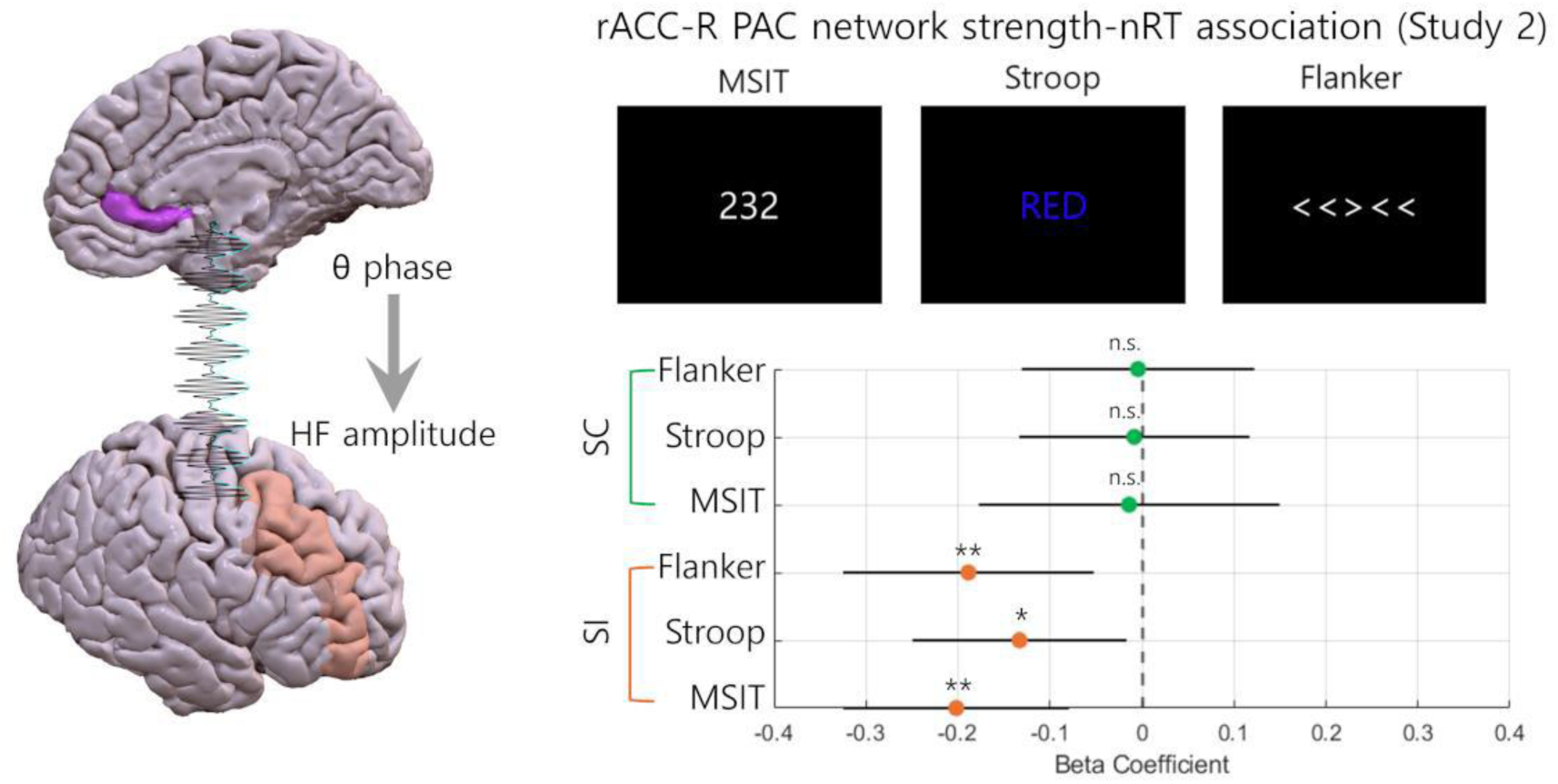
Replication and cross-task generalization of the rACC-R theta-PAC flexibility signal (Dataset 2; intracranial EEG). Using the same analysis pipeline applied in Dataset 1, an rACC-R theta-centered PAC network (PC1) was derived separately for the MSIT, Stroop, and Flanker tasks in Dataset 2. Across all three tasks, stronger rACC-R PAC network coupling was associated with faster nRT on SI trials, but not on SC trials, indicating that the SI-specific negative PAC–behavior relationship generalized across tasks and across an independent cohort. Points indicate beta coefficients from LMER and horizontal lines indicate 95% confidence intervals. Asterisks indicate FDR-corrected significance. * p<0.05, ** p<0.01, *** p<0.001.

### Pre-trial dACC-R PAC predicts subsequent flexible adjustment

Up to this point, we have described a within-trial, reactive mechanism whereby ACC-centered PAC supports adjustment after control-demand change. As an exploratory analysis, we next asked whether a pre-trial ACC signal might also predict subsequent flexible adjustment on the upcoming trial, building on earlier work showing that pre-trial PAC can predict faster conflict adaptation ^43^. Specifically, we tested whether pre-trial ACC theta–HFO PAC was associated with subsequent performance in a manner analogous to the reactive flexibility signal. We examined theta–HFO PAC originating from four ACC subregions (dACC-L, dACC-R, rACC-L, rACC-R) during the interval before participants encountered the upcoming control demand. We broadened this anatomic window beyond rACC-R to account for the possibility that pre-trial adjustment-related signals might rely on partially distinct ACC subcircuits.

We hypothesized that a pre-trial signal supporting subsequent flexible adjustment would predict faster nRT specifically when the upcoming trial required a change in control state (that is, on SI trials), and not when the upcoming trial repeated the prior control state (SC trials). Consistent with this prediction, pre-trial dACC-R–dlPFC-L theta–LG PAC was associated with faster nRT on subsequent SI trials (coefficient = – 0.05, *t* = –2.86, *p*FDR = 0.023; FDR-corrected across 4 × 10 theta–LG PAC pairs; Supplementary Figure 7), but not on SC trials. No other ACC-centered PAC, including rACC-R pairs, showed a significant effect under either condition.

These findings suggest that pre-trial theta-phase coordination associated with subsequent flexible adjustment engages the more dorsal ACC, whereas within-trial reactive adjustment is more strongly linked to the rostral ACC, echoing previous reports of a functional dissociation between these ACC subregions ^3^. In computational terms, this pattern is consistent with a distinction between a pre-trial signal that prepares for upcoming adjustment and a within-trial signal that supports reconfiguration once control-demand mismatch is encountered.

### Stimulation of Internal Capsule Enhances Control Flexibility (Dataset 1)

We next investigated whether internal capsule stimulation (ICS) could enhance control flexibility, based on past work showing effects of ICS on cognitive control task performance ^28,29,44^. In Dataset 1, 6 of the 21 underwent ICS during the MSIT ^28^. Each patient performed the task in both Stim ON (stimulated blocks, ICS randomly in 50% of trials) and Stim OFF conditions (all blocks from Stim OFF sessions and the first block from the Stim ON session prior to stimulation effects; Figure 4A).

**Figure 4.**
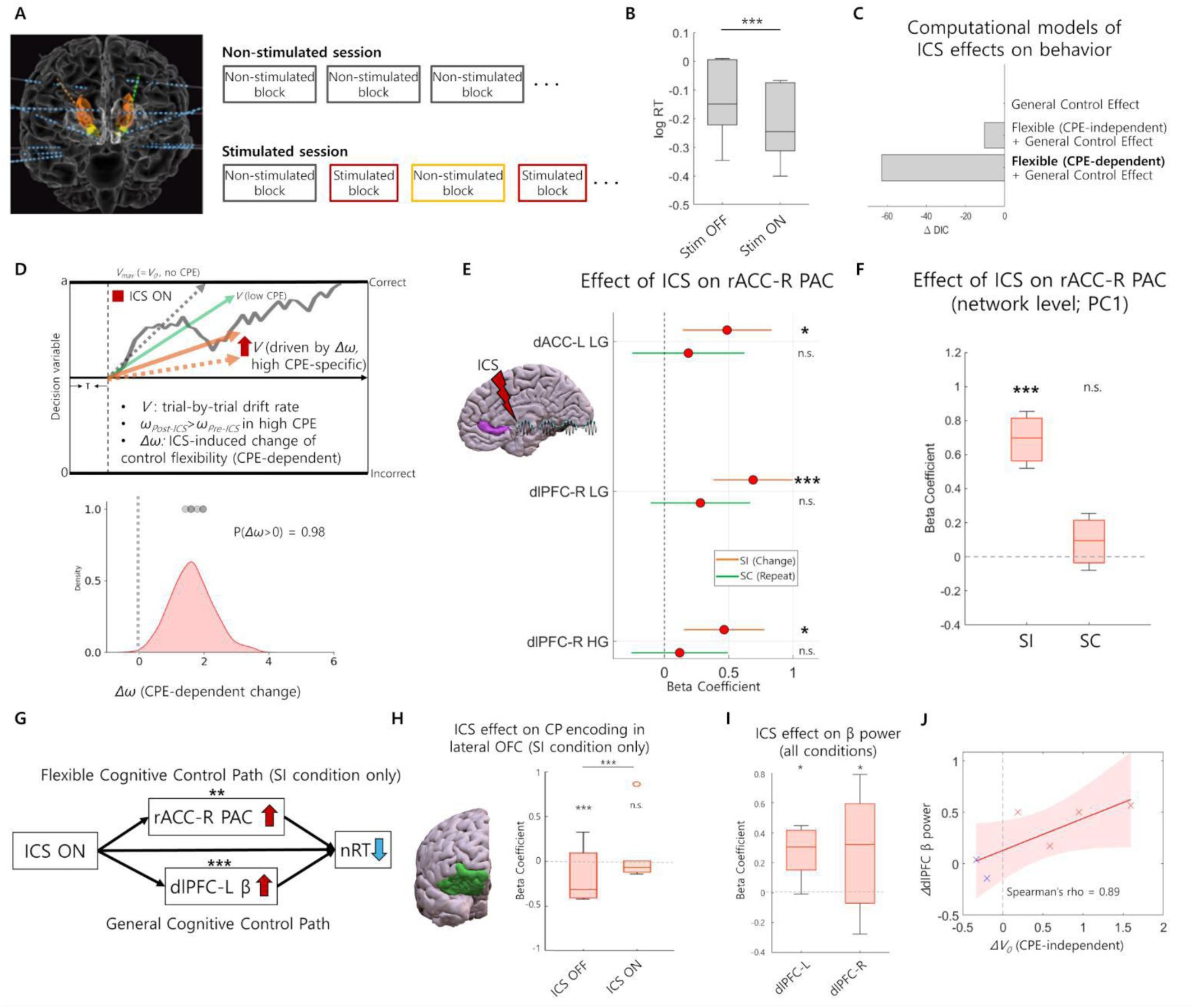
Internal capsule stimulation (ICS) enhances flexible adjustment via rACC-R theta PAC modulation (Dataset 1; iEEG). (A) Experimental design: six of the twenty-one patients performed the MSIT during both Stim OFF and Stim ON blocks. (B) ICS significantly decreased log RT overall. (C) Comparison of computational models of ICS effects on behavior. Among three modified adaptive DDMs, the model combining a general-control component with a CPE-dependent flexibility component provided the best fit. (D) Schematic of the flexibility parameter change, Δω, in the adaptive DDM and the group posterior distribution showing that ICS increased flexibility under high-CPE conditions. (E) ICS increased rACC-R theta-centered PAC in the three key PACs—rACC-R theta–dACC-L LG, rACC-R theta–dlPFC-R LG, and rACC-R theta–dlPFC-R HG—on SI trials but not SC trials. (F) At the network level, ICS increased the rACC-R PAC network (PC1) on SI trials but not SC trials. (G) Multilevel mediation analysis revealed that the rACC-R theta PAC increase partially mediated ICS-related nRT speeding under control-state change (SI), while bilateral dlPFC beta power, a general control signal, partially mediated a change-independent nRT reduction (SC). (H) ICS also suppressed interference from conflict-history (CP) encoding in the right lOFC during high-CPE trials, further supporting enhanced flexibility. (I) ICS increased bilateral local non– phase-locked beta power, a signal associated with general cognitive control (*Supplementary Methods & Results*). (J) ICS-induced increases in dlPFC beta power were associated with ΔV_0_, which indexes change in general cognitive control. Boxplots show median and interquartile range; whiskers denote the data range. In forest plots, circles indicate beta coefficients and horizontal lines indicate 95% confidence intervals. Asterisks indicate frequentist significance where applicable. * p<0.05, ** p<0.01, *** p<0.001.

In prior work ^28^, ICS significantly reduced MSIT RT across both conflict and non-conflict conditions (p<0.001, LMER; Figure 4B), and we showed that this arose from improved evidence/conflict processing ^45^. Here, we asked whether that RT improvement reflected only a more general increase in cognitive control, or whether it also included an enhancement of flexible adjustment when there is a control demand change.

To address this question, we compared three modified adaptive drift-diffusion models (adaptive DDMs) that differed in how ICS affected decision-making parameters: (1) a general-control model, in which ICS increased baseline drift rate (V₀) only; (2) a general-control plus CPE-independent flexibility model, in which ICS increased both V₀ and the flexibility parameter ω irrespective of mismatch level (CPE); and (3) a general-control plus CPE-dependent flexibility model, in which ICS increased ω proportionally with trial-wise CPE. The third model provided the best fit, demonstrating that ICS enhances CPE resolution specifically during high-CPE conditions (context-dependent flexibility enhancement; ΔDIC∼= −52 with the second best model, Figure 4C). Specifically, the group-level posterior for Δω indicated that ICS increased flexibility under high-CPE conditions (p(Δω > 0) = 0.982; observed in 6/6 participants; Figure 4D), whereas the general-control parameter ΔV₀ increased in 4/6 participants, with a weaker group-level posterior (p(ΔV₀ > 0) = 0.827). Thus, although ICS had a main effect on overall RT, the best-fitting model indicated that this effect was not exhausted by a context-independent speeding effect and instead included a distinct enhancement of flexibility when control-demand mismatch was high. This interpretation also aligns with prior work showing that stimulation can increase drift-rate-related evidence processing ^45^, while extending that account to include a mismatch-sensitive flexibility component.

We next asked whether ICS modulated the same rACC-R-centered PAC mechanism identified in the stimulation-OFF analyses. Because our main results showed that stronger rACC-R PAC was associated with faster performance specifically when control demand changed from the previous trial, we hypothesized that ICS would enhance this same PAC signal selectively under those conditions, thereby improving flexible adjustment. We therefore tested whether ICS increased rACC-R theta-centered PAC more strongly on SI than on SC trials. Focusing on the 18 PAC pairs that showed flexibility-related negative SI × nRT interactions in the preceding section, we found that ICS selectively increased PAC between rACC-R theta phase and HFO amplitude in key regions including dlPFC-R LG/HG and dACC-L LG on SI trials (all pFDR < 0.05; Figure 4E), but not on SC trials (all pFDR > 0.05). At the network level, ICS likewise increased coupling within the rACC-R PAC network specifically on SI trials (coefficient = 0.60, *t* = 3.76, *p* < 0.001; Figure 4F), with no effect on SC trials. Thus, ICS enhanced the same rACC-R-centered PAC configuration that was associated with faster performance when control demand changed.

We also asked whether ICS might facilitate adjustment by suppressing history-related signals that could interfere with current performance. In the stimulation-OFF condition, a right lateral OFC LG cluster encoded conflict probability (CP), reflecting carryover of recent conflict history. ICS abolished this CP encoding selectively on SI trials (ICS × CP interaction: *t* = 3.98, coefficient = 0.29, pFDR < 0.001, corrected across 10 CP-encoding clusters; Figure 4H), but not on SC trials (pFDR = 0.803). This pattern suggests that ICS may promote flexible adjustment not only by enhancing rACC-R-centered PAC, but also by reducing the influence of prior conflict history when current control demand changes.

Beyond this flexibility-specific pathway, ICS also enhanced a more general-control neural signal. Bilateral dlPFC beta power increased with stimulation, mediated a condition-independent decrease in nRT, and correlated with the behavioral marker of general-control improvement (ΔV₀; Figure 4G, I, J). Thus, Dataset 1 suggests a dual effect of ICS on cognitive control: a flexibility-related effect expressed when control demand changes and linked to rACC-R-centered PAC, and a more general effect on performance linked to local dlPFC beta power.

Together, these findings indicate that ICS does not merely speed behavior globally. Instead, stimulation appears to enhance flexible adjustment through a dissociable rACC-R-centered PAC mechanism that is selectively recruited when control demand changes, while also exerting a parallel general-control effect through dlPFC beta power. Clinical improvement (e.g., better control of obsessions/compulsions or negative mood regulation) may thus reflect enhanced flexibility and/or general information-processing capacity.

Because Dataset 1 did not include psychiatric patients, we examined this possibility in a separate clinical cohort with internal capsule DBS that modulated MSIT RTs (IC DBS; Dataset 3) ^29^.

### IC DBS in TRD Patients: Flexible Control Enhancement is associated with Clinical Response

In Dataset 3, a psychiatric cohort ^29^,chronic IC DBS significantly reduced RT during the MSIT without compromising accuracy, consistent with enhanced cognitive control. However, the magnitude of overall RT reduction did not predict clinical response ^29^. We therefore asked whether the clinically relevant component of DBS was a general improvement in cognitive control, or the more specific flexibility-related component identified in Dataset 1.

Given prior evidence linking cognitive flexibility to improved clinical outcomes across psychiatric disorders, including depression (e.g., utilization of coping skills to reduce distress;) ^46^ and OCD ^47^, we hypothesized that DBS-induced enhancement of flexible adjustment, rather than general cognitive control alone, would be more closely associated with treatment response. Using the adaptive DDM with ICS effects identified in Dataset 1 (Methods), we assessed DBS effects on the general-control parameter (ΔV₀) and the flexibility-related parameter (Δω). DBS significantly increased ΔV₀ at the group level (p(ΔV₀ > 0) = 0.985; Figure 5B, left), consistent with Dataset 1 and prior rodent findings. In contrast, Δω was not significantly increased at the group level in this dataset (p(Δω > 0) = 0.717; Figure 5B, right), indicating that not all patients showed improved control flexibility under DBS.

**Figure 5.**
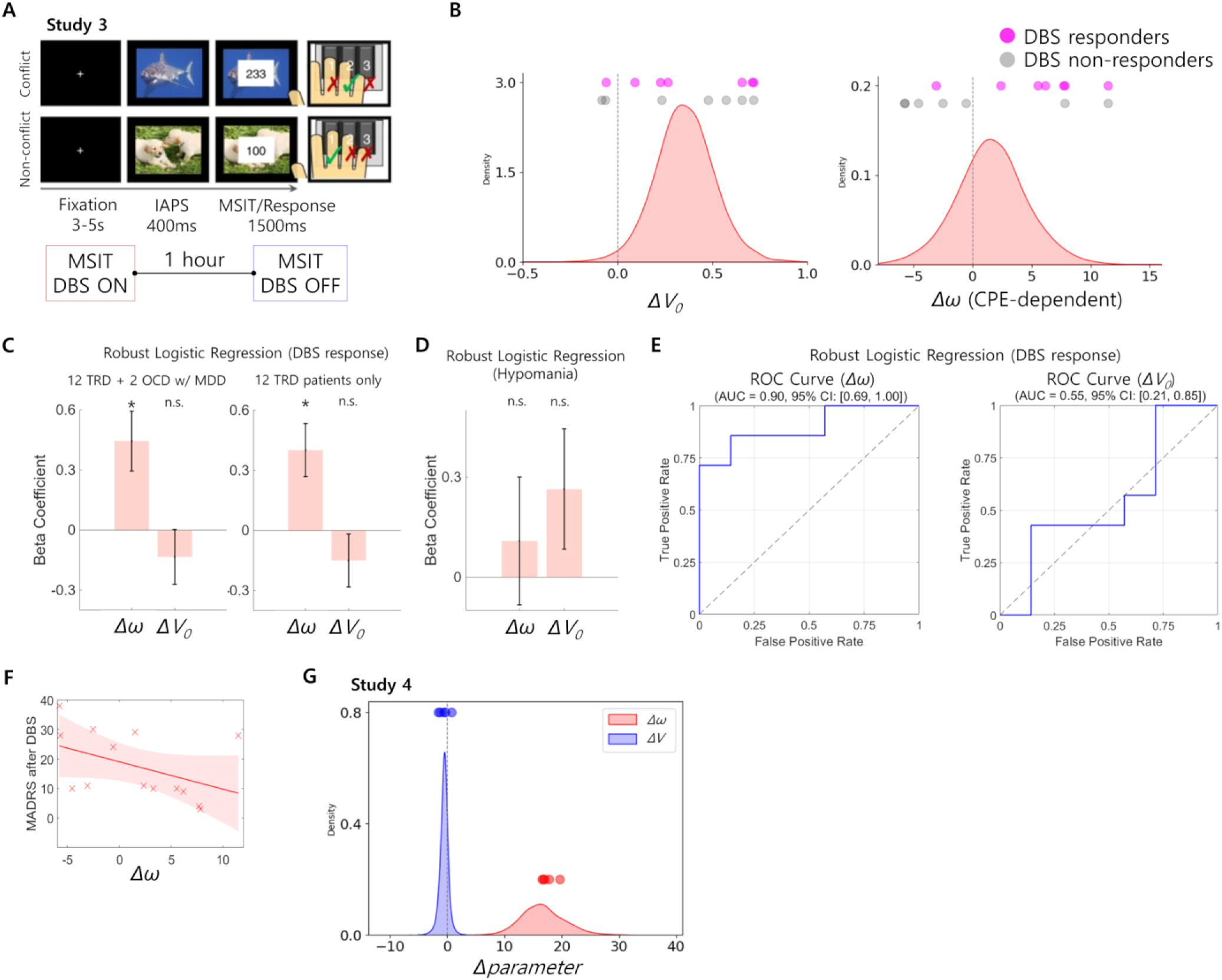
Clinical relevance and replication of flexibility-related control enhancement during IC DBS (Dataset 3 and 4). (A) Experimental design of Dataset 3: Twelve treatment-resistant depression (TRD) patients and two treatment-resistant OCD patients performed the MSIT task during both Stim OFF and Stim ON blocks. (B) Posterior distributions of IC DBS-induced changes in the general-control parameter (ΔV₀) and the flexibility-related parameter (Δω), with individual participants shown above each distribution and colored by clinical response status. Adaptive DDM revealed a non-specific increase in the ΔV₀ following DBS in both responders (6/7) and non-responders (5/7), while an increase in the Δω—reflecting improved flexibility under high-CPE (SI) conditions—was observed in the majority of responders (5/7) but in fewer non-responders (2/7). (C) Robust logistic-regression coefficients showing that Δω, but not ΔV₀, was associated with clinical response in the total cohort (N=14) and in the TRD-only subsample (N=12). (D) Neither Δω nor ΔV₀ was associated with hypomania. (E) Receiver operating characteristic (ROC) analysis confirmed that Δω was reliably associated with treatment response (AUC = 0.90), whereas ΔV₀ was not. (F) Greater Δω was associated with lower post-DBS depressive symptom severity. (G). Dataset 4 analysis in a separate TR-OCD cohort including five TR-OCD patients with chronic VCVS/IC DBS implants. Group-level posterior distributions of DBS-induced change in the flexibility-related parameter Δω and the general-control parameter ΔV₀ are shown for clinically selected IC DBS settings compared with DBS OFF. Dots indicate participant-level posterior mean estimates. Clinically selected IC DBS settings increased Δω across all five patients, whereas ΔV₀ did not show a clear group-level increase. Together, these findings indicate that the clinically relevant behavioral effect of internal capsule DBS is better captured by enhancement of flexible adjustment than by a more general improvement in cognitive control. Asterisks indicate frequentist significance where applicable. * p<0.05, ** p<0.01, *** p<0.001.

Despite the absence of a group-level Δω increase, individual differences in DBS-induced flexibility enhancement were strongly associated with clinical response. In robust logistic regression including both Δω and ΔV₀ as predictors and sex as a covariate, Δω was significantly associated with clinical response in the full cohort (N = 14; coefficient = 0.44, p = 0.014; Figure 5C), whereas ΔV₀ was not (p = 0.348). This relationship remained significant in the TRD-only subgroup (Δω: coefficient = 0.43, p = 0.022; ΔV₀: p = 0.286). Neither parameter was associated with hypomania emergence (all p > 0.1; Figure 5D). Consistent with this pattern, most DBS responders showed increased Δω (5/7), whereas fewer non-responders did so (2/7; Figure 5B, right). By contrast, ΔV₀ was increased in both responders (6/7) and non-responders (5/7), consistent with a more nonspecific effect on performance.

Receiver operating characteristic analysis further supported this dissociation. Δω was significantly associated with treatment response (AUC = 0.90, 95% CI [0.69, 1.00], p = 0.005; Figure 5E), whereas ΔV₀ was not (AUC = 0.55, 95% CI [0.30, 0.81]). In addition, Δω was significantly associated with improvement in depressive symptoms (Spearman’s ρ = –0.58, p = 0.028; Figure 5F). Thus, although DBS enhanced a general-control parameter at the group level, the flexibility-related component of behavioral improvement was the aspect most closely associated with therapeutic benefit.

Together with Dataset 1, these findings indicate that the clinically relevant effect of IC DBS is better captured by enhancement of flexible adjustment than by a more general improvement in cognitive control. Although Dataset 3 did not include neural recordings, the behavioral association with Δω is consistent with the flexibility-related mechanism identified in Dataset 1, in which rACC-R-centered PAC tracked and mediated stimulation-related improvement when control demand changed.

### Clinically selected IC DBS settings enhance flexibility-related control in TR-OCD

To test whether the flexibility-related stimulation effect generalized to a more clinically established OCD DBS context, we analyzed five patients with treatment-resistant OCD who participated in an ongoing study of the cognitive effects of IC DBS ^48^ (IRB: STUDY00021264; see Methods). All patients had chronic IC DBS implants and completed the MSIT during DBS OFF and during multiple DBS ON settings. Because IC DBS for chronic, severe, TR-OCD has an FDA Humanitarian Device Exemption indication, this cohort provided an opportunity to test whether clinically selected IC DBS settings engage the flexibility-related component of control in a disorder prominently characterized by maladaptive rigidity and compulsive behavior. For each patient, we selected the DBS ON setting that most closely matched that patient’s chronic clinical configuration (clinically effective in 4/5 patients)and compared that setting with DBS OFF.

Consistent with prior work showing that internal capsule stimulation can improve cognitive-control task performance, clinically selected IC DBS settings reduced log RT overall (LMER; coefficient = −0.05, *t* = −7.74, p < 0.001). We then applied the same adaptive DDM framework used above to determine whether this behavioral effect was better characterized as enhancement of general control or flexibility-related control. In this independent clinical cohort, clinically selected IC DBS increased the flexibility-related parameter Δω at the group level (p(Δω > 0) = 1.00; Figure 5G), with increases observed across all five patients. In contrast, the general-control parameter ΔV₀ did not show a clear group-level increase (p(ΔV₀ > 0) = 0.173; Figure 5G).

Thus, in a separate TR-OCD cohort, clinically selected IC DBS settings preferentially enhanced the flexibility-related component of control, providing convergent support for the idea that clinically optimized IC DBS may enhance flexible adjustment rather than exerting only a nonspecific general-control effect.

## Discussion

Across two independent intracranial datasets, we identified right rostral ACC (rACC-R) theta-centered PAC, particularly theta–gamma coupling with dlPFC-R and dACC-L, as a neural signature of efficient adjustment when control demand changes from one trial to the next. In Dataset 1, ICS enhanced this same adjustment-related signal selectively under conditions requiring control-state reconfiguration. In Dataset 3, the flexibility-related behavioral component of DBS effects, rather than the more general control component, was associated with clinical response. In Dataset 4, clinically effective IC DBS settings in a separate TR-OCD cohort preferentially enhanced the same flexibility-related component. Together, these findings support a model in which rACC-R theta-centered network coordination contributes to rapid control-state reconfiguration and suggest that this flexibility-related process may be the clinically relevant aspect of internal capsule neuromodulation in treatment-resistant psychiatric illness. The right-sided predominance aligns with other recent work suggesting that right-sided brain stimulation is more effective in enhancing cognitive control ^28,49^, and the emphasis on ACC and dlPFC aligns with a recent study on tracts mediating cognitive control improvement ^48^.

These findings fit naturally within contemporary models of cognitive control ^7,12,13,32,50^. Broadly, those models propose that the ACC detects when current control settings are no longer appropriate and helps coordinate reallocation of control, whereas LPFC maintains and implements task-relevant rules ^7,15,51,52^. Several theoretical and animal studies further suggest that this coordination is implemented at the network level through oscillatory interactions, particularly theta-phase-based modulation of higher-frequency activity in downstream executive regions ^15^. Primate work has shown that ACC–LPFC theta–gamma PAC accompanies attentional shifting ^17^, and recent rodent studies have further clarified how ACC signals can reshape frontal activity during flexible switching ^9,11^. In particular, Lam et al. provided evidence that ACC error-monitoring signals are transmitted through thalamus to promote PFC reconfiguration during flexible switching ^,11^. In that context, our main contribution is to identify a human intracranial signal consistent with that framework: rACC-R theta-HFO PAC, particularly with dlPFC and dACC gamma, that is selectively engaged when control demand changes and is associated with faster performance under those conditions, suggesting a role in efficient control-state reconfiguration.

The present findings also extend prior human neurocomputational work on adaptive control. Jiang et al. proposed that cognitive control is adjusted through trial-by-trial prediction of changing control demand, with an unsigned prediction error of congruency (control prediction error) resolution associated with flexible control adjustments ^3^. In that study, stronger rostral ACC encoding of this unsigned CPE was associated with less RT slowing, whereas dACC and dlPFC were linked to predicted control demand and control implementation. Our results align well with that framework while adding a network-physiological mechanism that was previously missing. Specifically, the most reproducible effects involved rACC-R theta-centered PAC with dlPFC-R and dACC-L gamma, and these same PACs generalized from the task-level SI contrast to model-derived unsigned CPE. Our adaptive DDM and model-derived CPE analyses further suggest that this adjustment-related rACC-centered PAC signal scales with the magnitude of control-demand mismatch (unsigned CPE). Thus, the current study complements Jiang et al. by showing that the human adjustment signal is expressed not only in rostral ACC activity at the level of fMRI, but also as an rACC theta-centered oscillatory coordination signal linking the principal hubs of the cognitive control network.

This broader interpretation also fits computational theories of ACC that frame its role in terms of valence-neutral mismatch or surprise signals that drive control adjustment ^10,12,13,32^. In the Predicted Response-Outcome (PRO) model and related work, ACC is proposed to signal deviations between expected and observed events, including unexpected occurrences and unexpected non-occurrences, in a valence-neutral manner ^12,13^; recent work further suggests that such ACC surprise signals can drive both inhibitory and motivated control ^10^. Consistent with that view, the rACC-R PAC–nRT correlation was specific to control-state change conditions, which were better captured by unsigned than signed formulations of mismatch/surprise (SI and CPE). This mismatch-specific effect was present across both directions of control-state change, thus suggesting that rACC-R PAC is a network mechanism for flexible adjustment after control-demand mismatch, in a manner similar to that proposed in the PRO model.

We also observed a dissociation between within-trial and pre-trial ACC signals. The within-trial flexibility effect was centered on rACC-R theta–gamma PAC with dlPFC-R and dACC-L, whereas the pre-trial effect involved dACC-R–dlPFC-L theta–LG PAC. We interpret this distinction cautiously. Because the pre-trial signal preceded the upcoming trial, it occurred before the system could know whether a control-demand mismatch would actually occur, making it difficult to conclude that it reflects proactive control in the strongest theoretical sense ^53^. A more conservative interpretation is that pre-trial dACC-centered coordination reflects a preparatory state that is associated with better subsequent adjustment when the upcoming trial requires control-state reconfiguration, whereas within-trial rACC-R-centered PAC is more directly linked to adjustment once mismatch is encountered. This pattern is broadly similar to Jiang et al., who reported that dorsal ACC and dlPFC were linked to predicted or proactive control, whereas rostral ACC was linked to reactive control associated with the resolution of unexpected residual conflict ^3^.

Neuromodulation techniques such as repetitive transcranial magnetic stimulation and DBS can modulate dysfunctional circuits in treatment-resistant psychiatric illness ^54^. Internal capsule (IC) stimulation, in particular, has shown benefit in disorders marked by cognitive rigidity, including OCD, depression, and addiction ^21–23,55^, although the underlying mechanisms remain incompletely understood. One hypothesis is that IC DBS improves cognition by synchronizing key control regions such as ACC and LPFC ^29^. Consistent with this idea, prior studies showed that IC stimulation enhances cognitive control and is accompanied by increased medial and lateral prefrontal theta activity ^28,29^. Cross-species work further showed that rodent mid-striatal stimulation, an IC-related manipulation, reduced response times in a set-shifting task, and computational modeling across rodents and humans linked these benefits primarily to increased drift rates without loss of accuracy ^30^. Because these previously reported effects generalized across task conditions, they were most consistent with a broad enhancement of cognitive control rather than a context-specific improvement in flexibility.

Our results extend that account in two important ways. First, the adaptive DDM showed that the behavioral effect of ICS was not limited to a general increase in drift rate, but was best explained by adding a separable flexibility-related component that emerged when control-demand mismatch was high. Second, ICS selectively strengthened the same rACC-R theta-centered PAC mechanism that, in the stimulation-OFF analyses, was associated with faster performance when control demand changed. In particular, ICS increased rACC-R theta–gamma PAC with key cognitive control regions, including dlPFC and dACC, specifically on trials requiring control-state reconfiguration, and this enhancement partially mediated the stimulation-related RT benefit. These findings suggest that rACC-R theta-centered PAC is not merely correlated with flexible adjustment, but is one mechanism through which IC stimulation enhances it. Relatedly, ICS also reduced OFC encoding of conflict history on trials requiring control-state reconfiguration, which may help the system shift away from outdated control-state representations when current demands change. At the same time, ICS independently increased dlPFC beta power, consistent with a parallel neural pathway for more general cognitive control improvement. Overall, these results suggest that IC stimulation has dual effects on cognition: a broad general-control effect, consistent with prior work, and a distinct flexibility-related effect that is expressed specifically when control-state reconfiguration is required.

Datasets 3 and 4 suggest that this flexibility-related component may be particularly relevant to clinical response following internal capsule DBS. Although DBS increased the general-control parameter at the group level, only the flexibility-related parameter was associated with clinical response and symptom improvement. These results raise the possibility that, in depression, flexible adjustment after control-demand mismatch may serve as a clinically meaningful behavioral marker of response. This interpretation also fits with emerging work in other psychiatric populations showing altered neural tracking of unsigned, control-related surprise: methamphetamine-dependent individuals showed reduced neural recruitment for a Bayesian surprise signal linked to adaptive inhibitory adjustment ^56^, and women remitted from bulimia nervosa showed attenuated control-related unsigned prediction-error signaling under specific internal-state conditions ^57^. More broadly, these findings raise the possibility that impaired tracking of control-demand mismatch and flexible adjustment to it may represent a clinically meaningful dimension across disorders marked by maladaptive inflexibility. Given the simplicity of the MSIT task, flexibility measures could serve as practical biomarkers during DBS programming ^58^.At the same time, the Dataset 3 clinical analyses should be interpreted cautiously. Dataset 3 did not include neural recordings, and the relationship between flexibility enhancement and symptom improvement was cross-sectional. Dataset 4 extends this clinical interpretation to a separate TR-OCD cohort with chronic IC DBS. Clinically selected DBS settings that generally improved OCD also increased Δω across all five patients, whereas ΔV₀ did not show a clear group-level increase. This suggests that IC stimulation may preferentially engage flexibility-related control across disorders marked by maladaptive rigidity.

A question remains: why did Dataset 1 and Dataset 4 show group-level flexibility-related enhancement, whereas Dataset 3 did not? One possibility is heterogeneity in stimulation targeting and programming context. Dataset 1 used experimentally controlled stimulation, and patients in Dataset 4 were programmed with awareness of IC anatomy and a desire to increase flexibility by engaging ACC and dlPFC (i.e., with awareness of the results from Datasets 1 and 3). Further, all patients in Dataset 4 were programmed by a single clinician with deep expertise in IC DBS. In contrast, patients in Dataset 3 were programmed by multiple clinicians, largely in the context of a study that favored stimulation in the more ventral capsule ^59^. Thus, the DBS settings in Dataset 3 engaged capsule subcomponents more heterogeneously. Conversely, this allowed clearer demonstration of the link between individual Δω changes and clinical response. The Dataset 3 result is also consistent with Dataset 1 findings where rACC-R PAC mediated flexibility gains. Thus, rACC-R PAC could serve as a neural biomarker for treatment response, alongside behavioral markers like enhanced control flexibility. These biomarkers may guide closed-loop DBS, delivering stimulation when flexibility or rACC PAC is low ^28^. Prior work has highlighted the potential utility of such biomarkers for optimizing DBS in depression ^60^ and OCD ^61^. However, we lacked neural recordings in Dataset 3 and 4 to directly support this mechanism, and the observed association between flexibility and symptom improvement was cross-sectional. Future studies are needed to test whether rACC-R PAC causally mediates clinical outcomes via improved flexibility.

Several limitations warrant caution. Sample sizes were modest, particularly for the ICS subset in Dataset 1 and the replication subset with rACC-R coverage in Dataset 2. Datasets 1 and 2 involved epilepsy patients, which may limit generalizability to healthy or psychiatric populations. The SI versus SC framework, while well suited to the present neural question, is related to but not identical with the classical Gratton framework; although our supplementary analyses showed that the main PAC effect was not reducible to a single transition direction or to a classical direction-specific adaptation pattern, future work using designs optimized to separate adaptive control from stimulus/response repetition effects would be valuable ^39^.

Although the adaptive DDM and prior ACC-focused control theories support an unsigned mismatch interpretation, alternative accounts of sequential effects based on carryover of prior-trial control states or event-file binding/unbinding effects involving conjunctive stimulus–response–rule representations may also contribute ^62^. The PAC findings also involve some analytic flexibility, because the precise rACC-R-centered topology and high-frequency targets emerged from a predefined search rather than from a single prespecified PAC pair. We mitigated this concern by applying the same model across the full 10 × 10 regional PAC space with multiple-comparison correction, validating the Dataset 1 finding across complementary PAC pipelines, and testing whether the resulting rACC-R-centered pattern generalized in an independent Dataset 2 rather than repeating the full discovery search. In addition, Dataset 3 and 4 lacked intracranial neural recordings, preventing direct testing of whether rACC-R-centered PAC mediates clinical improvement. Dataset 4 was small and did not include a formal responder-versus-nonresponder analysis, because 4 of the 5 patients were responders. It should thus be interpreted as evidence of flexibility-related target engagement rather than proof that Δω predicts clinical response. Finally, because the clinical analyses were cross-sectional, the observed associations should not be interpreted prospectively.

In conclusion, our results identify rACC-R theta-centered PAC, particularly coupling with dlPFC and dACC gamma, as a candidate human network mechanism for efficient adjustment when control demand changes. Internal capsule stimulation enhanced this same flexibility-related mechanism in Dataset 1, the corresponding behavioral component was associated with clinical response in Dataset 3, and clinically selected IC DBS settings preferentially enhanced this component in Dataset 4. Together, these findings advance a circuit-level account of flexible cognitive control and suggest that flexibility-related behavioral and neural signals may be useful targets for future neuromodulation strategies in depression, OCD, and other disorders marked by cognitive rigidity.

## Methods

### Ethics statement

All analyses of previously published datasets (Dataset 1-3) were conducted using de-identified data. Regulatory approvals and informed-consent procedures for the original data collections are described in the source publications for each dataset. For Dataset 4, the study was approved by the University of Minnesota Institutional Review Board (IRB: STUDY00021264). Participants voluntarily participated in the study and provided written informed consent prior to study procedures. All study procedures were conducted in accordance with the Declaration of Helsinki.

### Participants and Datasets

**Dataset 1 (Discovery dataset** ^28^): 21 adults (12 female) with medication-refractory epilepsy and no primary psychiatric diagnoses were recruited from inpatient epilepsy monitoring units. All underwent intracranial electrode implantation for clinical seizure localization and volunteered for research after electrode placements were finalized. Each participant had 8–18 depth electrodes placed bilaterally in key cortical and subcortical regions, including frontal, cingulate, and hippocampal areas. Post-operative CT scans were rigidly co-registered to the pre-implant T1-weighted MRI using FreeSurfer scripts (http://surfer.nmr.mgh.harvard.edu). Electrode contacts were manually identified ^63^ and automatically assigned anatomical labels with the Interactive Electrode Localization Utility ^64^, which applies a probabilistic atlas-based algorithm. These depth electrodes (with 8–16 platinum/iridium contacts) recorded at a 2 kHz sampling rate.

Participants performed the Multi-Source Interference Task (MSIT), a cognitive control task where they viewed three numbers (e.g., “2 2 1”) and were asked to press the number that differed from the other two within 1.75 seconds. Conflict trials involved a mismatch between the numeric identity and the spatial position of the correct response (e.g., “1” is in the 3rd position in “2 2 1”), requiring suppression of positional cues. Non-conflict trials had no such mismatch. Conflict and Non-conflict trials occurred randomly with rules preventing long runs of the same trial type (see Basu et al. ^28^ for details). This dataset was used to develop the analytic pipeline that ultimately tested whether phase–amplitude coupling (PAC) between the anterior cingulate cortex (ACC) and LPFC/hippocampus correlates with flexible adjustment after control-demand change to support cognitive flexibility in a non-psychiatric population.

In a subset of six patients, brief internal capsule stimulation (ICS) was applied during the MSIT. These patients performed task blocks with and without stimulation, allowing within-participant comparisons of ICS ON vs. OFF effects. In the ICS ON blocks, half the trials (randomly selected) received 600 ms stimulation trains (130 Hz, 2–4 mA), while the remaining trials were unstimulated. Only the non-stimulated trials from ICS ON blocks were used for LFP analyses to avoid stimulation artifacts. Stimulations were delivered to either dorsal or ventral internal capsule, and on either hemisphere, creating within-participant comparisons. Some patients also completed closed-loop stimulation blocks, but those were excluded from the primary ICS ON analyses due to potential confounds from behavior-triggered stimulation.

**Dataset 2 (Replication dataset** ^31^): This independent sample included 16 medication-refractory epilepsy patients undergoing iEEG monitoring via depth electrodes in frontal, parietal, and temporal regions. Depth-electrode contacts were localized with the iELVis workflow ^65^: the pre-implant T1 MRI was segmented in FreeSurfer, the post-implant CT was co-registered to that MRI, contacts were manually tagged in BioImage Suite ^66^, and each coordinate was automatically mapped to a Desikan–Killiany cortical parcel (or subcortical/white-matter label) by the FreeSurfer localization tool (http://surfer.nmr.mgh.harvard.edu); contacts falling in white matter, ventricles, cerebellum, or unlabeled tissue were excluded from analysis. Participants completed three cognitive control tasks—MSIT, Stroop, and Flanker—without any DBS. Each session consisted of 18 blocks (30 trials per block), covering all three tasks. In the Stroop and MSIT tasks, responses were given verbally (recorded at 8,192 Hz), while Flanker responses used key presses. These tasks all included Conflict trials requiring suppression of irrelevant information (e.g., semantic vs. color in Stroop, flanker arrows vs. center arrow in Flanker). This dataset allowed replication of neural findings from Dataset 1 in a separate, non-psychiatric cohort. The Dataset 2 analysis used the same preprocessing, SI/SC definitions, PAC computation, rACC-R PAC network construction, and LMER specification defined in Dataset 1, with no task-specific reselection of region-frequency effects.

**Dataset 3 (TRD/TR-OCD Cohort** ^29^): 14 adults (6 males, 8 females), ranging from their 30s to 70s, with severe, treatment-resistant psychiatric illness—12 with treatment-resistant major depressive disorder (TRD) and 2 with treatment-resistant obsessive–compulsive disorder (TR-OCD)—participated in this study. All had failed multiple prior interventions (e.g., medications, psychotherapy, ECT) and received chronic IC DBS implants as part of previous clinical trials (NCT00640133, NCT00837486, NCT00555698). Participants completed an affective variant of the MSIT task under both DBS OFF and DBS ON conditions. Clinical response was defined as a ≥50% reduction in MADRS scores (for MDD) or ≥35% reduction in YBOCS scores (for OCD). This dataset was used to assess whether DBS-induced improvements in flexible adjustment were associated with clinical outcomes in patients with severe mental illness.

**Dataset 4 (TR-OCD Cohort):** Dataset 4 included five patients (VCVS001–VCVS005) with treatment-resistant OCD and chronic VCVS/internal capsule DBS implants who participated in the VCVS Cog Study (IRB: STUDY00021264). This study was designed to evaluate the acute cognitive effects of DBS parameter settings during the MSIT. Participants completed the MSIT during DBS OFF and during multiple DBS ON stimulation settings (Supplementary Table 3). Each MSIT session included 100 trials, except for one session that included 102 trials. The number of DBS OFF sessions was 3, 4, 3, 3, and 3 for VCVS001–VCVS005, respectively. The number of DBS ON sessions across tested stimulation settings was 24, 24, 24, 20, and 24 for VCVS001–VCVS005, respectively, with each DBS ON session corresponding to a distinct stimulation setting. Across the stimulation survey, DBS was delivered at 130 Hz using monopolar stimulation of left and/or right internal capsule contacts. We tested each contact in monopolar mode, amplitudes of 2–4 mA, and pulse widths of 150–210 μs, with exact setting combinations varying slightly across participants based on protocol completion and clinical considerations.

4 of the 5 participants responded to IC DBS, either defined as a Yale-Brown Obsessive-Compulsive Scale reduction of ≥33% or a substantial improvement in quality of life (one patient who returned to and completed a challenging academic program, gained independent living skills, reduced restrictive eating, and increased socialization with peers, but did not numerically change their YBOCS). As such, this dataset could not be used to test correlations of model parameters with response, but could test whether clinically effective DBS changed those parameters.

### Behavior data analysis

Trials with RT < 100 ms were excluded from behavioral analyses as premature responses ^67,68^. For Dataset 1, this resulted in 6896 trials including both ICS ON and OFF conditions (21 patients; with 2.1% incorrect trials; 6751 correct trials). ICS OFF trials were 5976 trials (with 2.3% incorrect trials; 5838 correct trials) and ICS ON trials were 920 trials (6 patients; with 0.7% incorrect trials; 913 correct trials). For Dataset 2 there were only conditions without ICS/DBS. The number of trials included were 2558 for MSIT (16 patients; 0% of incorrect trials; 2558 correct trials), 2657 for Stroop (16 patients; 0.8% of incorrect trials; 2655 correct trials), 2724 for Flanker (16 patients; 4.3% of incorrect trials; 2606 correct trials). For the Dataset 3, there were 3999 trials in total including both DBS ON and OFF conditions (14 patients; with 1.7% of incorrect trials; 3930 correct trials). DBS OFF trials were 1988 trials (with 1.7% of incorrect trials; 1954 correct trials) and DBS ON trials were 2011 trials (6 patients; with 1.7% of incorrect trials; 1976 correct trials).

Given that the average accuracy of the task was very high (>95%), we focused on analyzing RT. We used log-transformed RT (log RT) as the primary behavioral measure, as it reduces the positive skew of the RT distribution and approximates a Gaussian distribution ^69^. Our prior work ^29,45^ achieved the same using a generalized linear model with a gamma distribution. We switched to the log-Gaussian model here to enable more computationally efficient fitting of multiple models during large-scale trial-wise neural analyses compared to a gamma GLMM. For the log RT analyses, we only utilized correct trials. First, we analysed the effects of conflict and SI at the trial level using a linear mixed effect regression (LMER). As in our earlier work ^28^, the block number was used as a covariate and the participant was used as a random effect variable as below (this was the same for all LMERs used in this study):

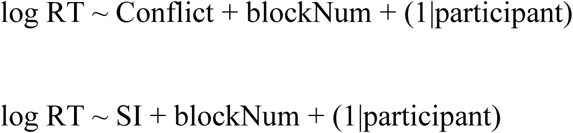

Here, Conflict refers to trials in which the target’s identity and spatial position are mismatched, requiring resolution of competing response tendencies. Sequential Incongruence (SI) denotes trials where the current trial type differs from the previous one (e.g., Conflict following Non-conflict or vice versa), thereby indexing a change in control demand and serving as a task-level proxy for higher control-demand mismatch.

### Computational Modeling: Adaptive Drift-Diffusion Model (DDM)

In the behavioral analyses, we showed that sequential incongruency (SI), which indexes a change in control demand relative to the previous trial, is associated with longer RTs. In this computational modeling section, we formally defined the magnitude of control-demand mismatch as control prediction error (CPE) and tested whether a model that incorporates this mismatch signal into the decision-making process (the adaptive DDM model) explains participants’ RTs better than a basic model without a CPE component (the basic DDM model ^70^).

**Basic DDM:** This is the standard drift-diffusion model (DDM) ^70^ for two-alternative forced choice (we adapted it to our 3-alternative MSIT by focusing on target vs. non-target processing). Key parameters include the drift rate (*v*), which reflects the speed of evidence accumulation (higher *v* means faster information processing toward the correct response), and the decision boundary (*a*), which reflects the amount of evidence required (a larger *a* yields more cautious, slower decisions). Non-decision time (*τ*) captures sensory–motor delays. In the basic DDM, these parameters are fixed across trials of a given condition. Further, this DDM does not account for control history or adjustments based on previous trials. It assumes performance is driven only by current stimulus evidence quality.

**Adaptive DDM:** We extended the DDM by incorporating an internal estimate of conflict or control demand that is updated each trial – effectively modeling the participant’s ongoing estimate of conflict probability (CP). In this model, on each trial *k*, the participant has a predicted control demand CP(*k*) (ranging 0 to 1, where 1 means they fully expect conflict). We implement a simple reinforcement learning rule: after each trial, the estimate is updated based on whether conflict occurred or not (e.g., increase CP after an incongruent trial, decrease it after a congruent trial, akin to a delta-rule with a learning rate parameter). The control prediction error (CPE) on trial *k* is then defined as the absolute difference between actual conflict *C*(*k*) (1 for incongruent, 0 for congruent) and the predicted CP(*k*):

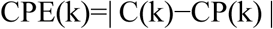

This term, which is an unsigned prediction error, represents surprise in control demand as in previous studies ^3,57^. We hypothesized that the drift rate *v* is modulated by CPE: if a trial’s conflict level is unexpectedly high, the drift rate (evidence accumulation) would be temporarily reduced due to the need to reallocate resources. We implemented this as:

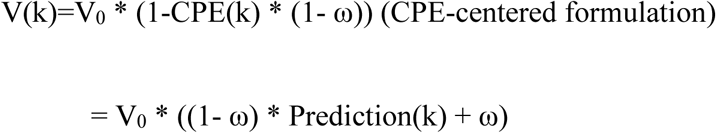

(Prediction-centered formulation; Prediction(k) means the predicted portion of the control demand, which means Prediction(k)=1-CPE(k))

Here, V₀ is the baseline drift rate (i.e., when no prediction error is present), and ω is a flexibility parameter (0 – 1) that captures an individual’s control flexibility given that a higher ω dampens the impact of a control–prediction error (CPE) on drift rate, allowing the agent to become more reactive to unexpected control changes. When ω is near 1, even a large CPE barely slows the drift, indicating robust, rapid adaptation. When ω is small, the same CPE markedly slows the drift, reflecting low flexibility and greater susceptibility to mismatch. We estimate logit ω (range –∞ to ∞) and transform it with a sigmoid so that ω lies within [0, 1]. All DDM parameters—baseline drift V₀, flexibility ω, and boundary separation a—were allowed to vary by stimulus condition (congruent vs. incongruent).

All models (basic DDM, adaptive DDM) were fitted to the trial-by-trial RT distributions and accuracy data using hierarchical Bayesian inference using HDDM package ^70^ (30,000 samples with 25,000 burn-in samples were drawn from four MCMC chains and the model convergence was assessed by testing whether the Gelman-Rubin statistic was <1.1) and we compared model fit using the Deviance Information Criterion (DIC), which penalizes model complexity. Lower DIC values indicate a better trade-off of fit vs. complexity.

Lastly, we conducted a posterior predictive check following the standard procedure in the HDDM package ^70^. For each participant and condition, we drew 500 parameter sets from the posterior distributions of the fitted parameters (V₀, ω, a, *τ*) and, for each set, simulated a complete trial–by–trial dataset that matched the original numbers of congruent and incongruent trials. This yielded 500 synthetic RT and accuracy distributions per participant. From each simulated dataset we calculated summary statistics—accuracy, median RT, and the 10th, 30th, 70th, and 90th RT percentiles—and then constructed 95 % credible intervals across simulations. The empirical statistics all fell within these intervals. We also computed Mahalanobis distances between the observed and predicted summary–statistic vectors; the consistently low values (Supplementary Table 1) provide additional evidence of satisfactory model fit.

**Effect of ICS on Adaptive DDM Parameters (Dataset 1 and 3):** In Dataset 1, we tested three alternative computational models to determine how ICS (DBS in Dataset 3) influences parameters within our adaptive drift-diffusion model (adaptive DDM). The best-fitting model (third model: CPE-dependent flexible control enhancement + general control enhancement) identified in Dataset 1 was applied to the analyses of Dataset 3.

The first model assumed that DBS affects only the general aspect of cognitive control by increasing the baseline drift rate, V_0_ (i.e., V_0__postICS=V_0_ + ΔV_0_). This effect was previously identified using a basic HDDM model. To further investigate whether ICS also modulates control flexibility, we examined two additional models that incorporate changes in the flexibility parameter ω (Δω).

In the second model (CPE-independent flexible control enhancement + general control enhancement), ICS is assumed to alter ω in a control-prediction error (CPE)–independent manner, defined as:

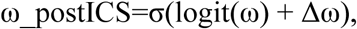

where σ(x)=1/(1 + exp(–x)) is the sigmoid function (used to constrain ω within the [0, 1] range), and Δω reflects the CPE-independent effect of ICS on the (logit-transformed) flexibility parameter.

In the third model (CPE-dependent flexible control enhancement + general control enhancement), ICS is assumed to modulate ω in a CPE-dependent manner, such that:

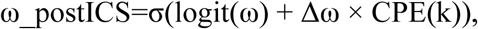

where Δω represents the magnitude of the ICS effect on logit(ω) that varies proportionally with the control prediction error on trial k.

### iEEG preprocessing

We utilized open iEEG datasets from previous studies (https://zenodo.org/record/5083120#.YOhvWehKiUk and https://zenodo.org/record/5085197#.YOhtouhKiUk for Dataset 1, pre-processed; https://klab.tch.harvard.edu/resources/XiaoEtAl_CognitiveControl.html for Dataset 2, pre-processed). For the details of pre-processing, please refer to Basu et al., ^28^ for Dataset 1 and Xiao et al., ^31^ for Dataset 2.

Briefly, for Dataset 1, signals were bipolar re-referenced, down-sampled to 1,000 Hz, and demeaned. Recordings were cleaned of 60 Hz line noise and its harmonics, and pathological channels with interictal epileptiform discharges (IEDs) were removed. For Dataset 2, line noise (60 Hz and its harmonics) were filtered out and the trials containing artefact (amplitudes larger > three standard deviations above the mean amplitude across all trials) were excluded from the analyses.

### Phase–Amplitude Coupling (PAC) Analysis

To assess trial-by-trial phase–amplitude coupling (PAC), we used a variant of the Modulation Index (MI) method originally introduced by Tort et al. ^14^, and adopted in recent studies ^40^. PAC was computed within a post-stimulus window of 0.1–1.4 s after stimulus onset following our previous work (Dataset 1 ^28^); or −0.5–0.5 s centered on the response in RT-locked analyses. For RT-locked PAC analyses, trials with RT < 500 ms were additionally excluded because PAC was computed from a 1 s segment centered on the response; including shorter-RT trials would cause the response-centered window to extend into pre-trial activity and yield truncated or non-comparable analysis segments. Time–frequency decomposition was performed using complex Morlet wavelet convolution from 2 to 150 Hz in logarithmic steps with 20ms time steps to extract the phase of low-frequency theta oscillations (4–8 Hz) and the amplitude envelope of high-frequency oscillations (beta (15-30 Hz), low-gamma (30-50 Hz; LG), and high-gamma (70-150 Hz; HG)). Data were epoched from −2 to +3 s relative to stimulus onset. Phase and amplitude estimates were obtained using 7-cycle Morlet wavelets to balance temporal and spectral resolution, and stimulus-locked PAC was quantified over the 0.1–1.4 s post-stimulus interval. For each electrode pair and trial, we binned the theta phase into 20 equal intervals and calculated the average high-frequency amplitude within each phase bin. The resulting amplitude–phase distribution was compared to a uniform distribution using Kullback–Leibler divergence to yield the Modulation Index (MI) ^14^, quantifying the strength of PAC.

Trial-wise MI values were then averaged across electrode pairs within 10 predefined anatomical regions of interest (ROIs) to generate ROI-level PAC features for subsequent analyses.

To relate PAC dynamics to behavior, we fitted a linear mixed-effects model for each of the 10 × 10 region pairs to determine whether the trial-by-trial association between PAC and normalized RT (nRT) differed depending on whether control demand changed or repeated. Normalized RT (nRT) was computed by z-scoring log RT separately within Conflict and Non-conflict trials. This normalization removed the large condition-level mean RT difference, allowing trial-level neural analyses to test whether activity tracked responses that were relatively faster or slower than expected for that condition, rather than simply recapitulating the conflict main effect. Participant-level differences in baseline speed were additionally accounted for in the mixed-effects models by including the participant as a random effect. The model included fixed effects for nRT, SI, their interaction, and block number, with participant as a random intercept:

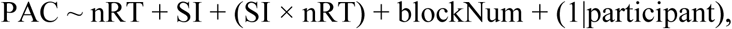

The nRT × SI interaction tested whether the PAC–nRT relationship was modulated when control demand changed from the previous trial (SI) relative to when it repeated (SC). P-values were FDR-corrected for 10 × 10 regional PAC pairs.

We next performed a post-hoc analysis on the 18 PACs that showed a significant negative SI × nRT interaction, testing PAC–nRT correlations separately within SI and SC trials using LMER. The LMER was conducted using the following equation for each condition:

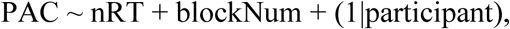

P-values were FDR-corrected across the 18 PAC pairs.

### Validation of Trial-by-Trial PAC Analysis Using Trial-Shuffling Methods

We validated the observed significant negative SI-related interaction effects on the nRT–rACC-R PAC correlation by restricting the analysis to electrode pairs whose trial-averaged PAC values were deemed significant based on a surrogate-based null distribution. Following a method used in a recent study ^40^, we generated 500 surrogate trial-averaged PAC values per channel by randomly pairing phase and amplitude signals from different trials (i.e., trial-shuffling). This procedure disrupts true phase–amplitude coupling while preserving the spectral content of each trial, thereby helping to rule out spurious PAC driven by signal autocorrelations or non-oscillatory transients. For each electrode pair, a z-score for the observed Modulation Index (MI) was calculated against this surrogate distribution. Pairs with z-scores exceeding 1.64 were classified as significant ^40^. We then repeated the same LMER analysis using only these significant pairs.

However, we note that our original PAC analyses included all electrode pairs; restricting the analysis to only “significant” pairs may omit condition-specific PAC fluctuations that emerge transiently in typically weaker couplings (e.g., during SI trials with fast RTs).

### rACC-R PAC Network Definition (PCA)

To summarize the multi-regional PAC findings, we performed a principal component analysis (PCA) on the trial-wise rACC-R PAC matrix (# of trials × # of connected region–frequency pairs). The first principal component (PC1) typically captured 78% of the variance and had roughly uniform positive weights across all region–frequency PAC pairs, thereby serving as an index of overall rACC-R-theta-phase-centered coupling strength on that trial (we termed this the rACC-R PAC network strength). PCA was performed separately for each dataset/task on the set of rACC-R-centered PAC pairs, and PC1 scores were used as a summary measure of network-level coupling. Because PC1 explained most of the variance and showed broadly similar positive loadings across PAC pairs, it likely reflects overall rACC-R-centered coupling strength within the predefined PAC set. We then used this network measure in statistical models relating PAC to RT including LMER that we mentioned above as well as quantile-averaged approach *(Supplementary Results*; Supplementary Figure 3).

### Effects of ICS on PAC

For the 18 PAC pairs (centered in rACC-R theta) that showed neural patterns of flexible adjustment in the stimulus-locked PAC analysis, we tested whether ICS increased rACC-R–centered PAC separately within SI and SC trials using LMER with the following equation:

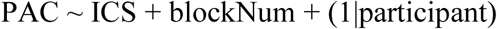

where ICS was coded as 1 (ICS ON) and 0 (ICS OFF). P-values were FDR-corrected across the 18 PACs. Only the non-stimulated trials from ICS ON blocks were used for LFP analyses to avoid stimulation artifacts.

We also tested whether ICS increased the rACC-R PAC network separately within SI and SC trials using the same LMER.

### Mediation analysis

The mediation analysis tests whether covariance between two variables (X and Y) can be explained by a mediator variable (M). Significant mediation is present when adding M to the model changes the X→Y slope, with the mediation effect quantified as a×b . We asked whether rACC-centered PAC (M) mediated the effect of ICS (X; on vs. off) on nRT (Y) during SI trials, that is, trials requiring control-state reconfiguration. Multilevel mediation analyses were fitted using the MATLAB MediationToolbox (Canlab), estimating the a path (X→M), the b path (M→Y), the c path (X→Y without M) and the mediation effect a×b (reduction of c path after inclusion of M). Inference used participant-clustered nonparametric bootstrapping (10,000 resamples). The primary model pooled stimulation sites; we then repeated the analysis separately for VS ICS and DS ICS.

### Analyses of Dataset 3

We fit the best adaptive DDM identified in Dataset 1 (Model 3: CPE-dependent flexible-control enhancement plus general-control enhancement) to the RT and accuracy data in Dataset 3. We examined the posterior probabilities of the DBS effect parameters, including ΔV_0_ (change of baseline drift; general control) and Δω (flexibility under high CPE).

We then used robust logistic regression (MATLAB function “robustfit”) to test whether changes in general control or control flexibility were associated with clinical response to DBS, using the model:

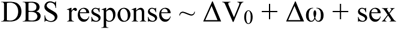

Analyses were conducted in the full cohort (N=14; 12 TRD, 2 OCD + MDD; 7 responders) and separately in the TRD subgroup (N=12; 6 responders). Sex was included as a covariate to account for potential sex-related variance in clinical response in this small mixed clinical cohort. To assess each parameter’s diagnostic value, we computed receiver operating characteristic (ROC) curves and area under the curve (AUC) from single-parameter models (DBS response ∼ parameter; separately for ΔV_0_ and Δω) using “perfcurve” in MATLAB. Uncertainty was quantified with 1,000 bootstrap resamples, and statistical significance was assessed with a 1,000-label permutation test; p-values reflect the proportion of shuffled AUCs exceeding the observed AUC. For the permutation test, clinical-response labels were randomly shuffled across participants while Δω and ΔV₀ values were held fixed. Finally, we tested associations between post-DBS MADRS scores and parameter changes using Spearman’s rho.

Lastly, we tested whether changes in general control or control flexibility were associated with hypomania using the model:

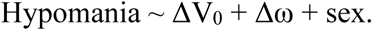

Hypomania was tested because it was a possible stimulation-emergent side effect after the IC DBS and could represent a nonspecific mood-elevation effect distinct from therapeutic response for depression ^71^.

### Analysis of Dataset 4

For the primary Dataset 4 analysis, we selected, for each patient, the DBS ON setting or set of settings that most closely matched that patient’s chronic clinical configuration. We compared those clinical-like settings with DBS OFF (Supplementary Table 4). This yielded 6, 12, 12, 6, and 9 clinical-like DBS ON sessions for VCVS001–VCVS005, respectively. When the chronic clinical configuration involved bipolar or multi-contact stimulation that was not exactly reproduced in the research stimulation survey, the closest available research setting was selected based on stimulated contact location and cathode involvement.

Behavioral effects were assessed using linear mixed-effects regression on correct-trial log RT:

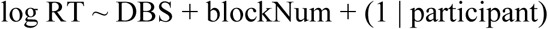

where DBS was coded as clinical-like DBS ON versus DBS OFF. We then applied the same adaptive DDM framework used for Datasets 1 and 3 to estimate DBS-induced change in the flexibility-related parameter Δω and the flexibility-independent general-control parameter ΔV₀.

## Acknowledgments

This work was supported by the MnDRIVE Brain Conditions initiative, the Minnesota Medical Discovery Team on Addictions, and the National Institutes of Health (NIH) under grants R01MH125429, R01NS120851, R01MH124687, R01MH123634, and UH3NS100548. We gratefully acknowledge the use of three publicly available datasets: Dataset 1 (https://zenodo.org/record/5083120 and https://zenodo.org/record/5085197), Dataset 2 (https://klab.tch.harvard.edu/resources/XiaoEtAl_CognitiveControl.html), and Dataset 3 (https://openneuro.org/datasets/ds001784). We thank the Kreiman Laboratory for generously sharing their dataset and related resources (Dataset 2). We also appreciate valuable discussions with Geoffrey Diehl, Evan Dastin-Van Rijn, Jeremiah Morrow, and Aaron McInnes, which helped improve the conceptual framing and interpretation of the results.

## Supplementary Materials

**Supplementary Figure 1.**
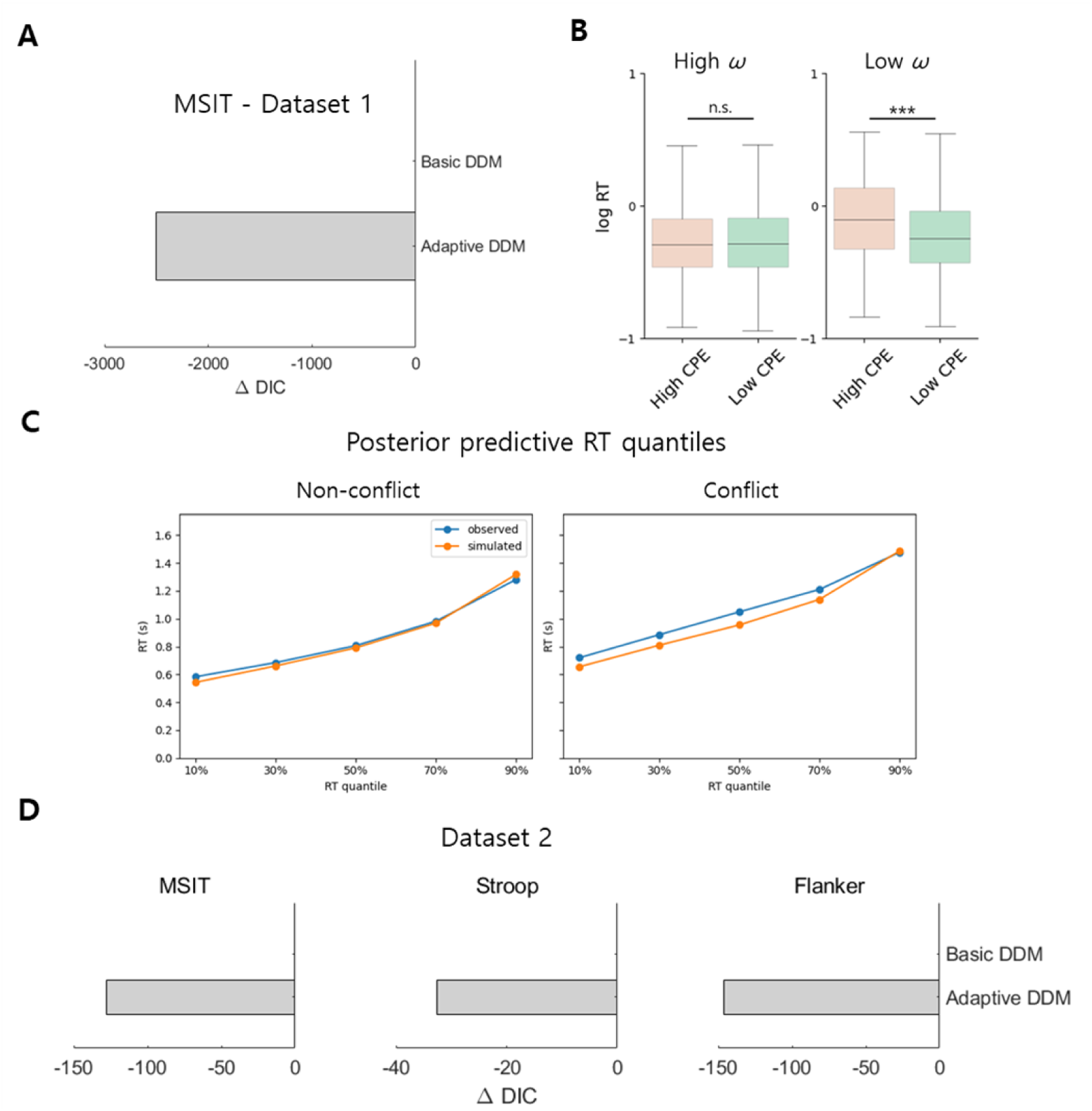
Computational modeling results. (A) In Dataset 1, fitting this adaptive DDM to trial-by-trial RT distributions and accuracy significantly outperformed a basic DDM (ΔDIC=–2502). (B) ω is a flexibility parameter (0 – 1) that captures an individual’s control flexibility given that a higher ω dampens the impact of a control–prediction error (CPE) on drift rate, allowing the agent to become more reactive to unexpected control changes. When ω is near 1 (left, high ω condition, ω=0.99, 10,000 simulations), even a large CPE barely slows the drift, indicating robust, rapid adaptation. When ω is small (right, low ω condition, ω=0.5, 10,000 simulations), the same CPE markedly slows the drift, reflecting low flexibility and greater susceptibility to mismatch. (C) Posterior-predictive RT quantiles for Dataset 1, shown separately for conflict and non-conflict trials to visualize condition-specific model fit. Observed and simulated RT quantiles showed reasonable agreement across conditions. (D) Model comparison results were replicated in the MSIT in Dataset 2 (ΔDIC=–127) and also held for other conflict tasks such as Stroop and Flanker (ΔDIC=–33 and –146, respectively), indicating that incorporating CPE better explains behavior than a fixed-drift rate model across studies. * p<0.05, ** p<0.01, *** p<0.001.

**Supplementary Figure 2.**
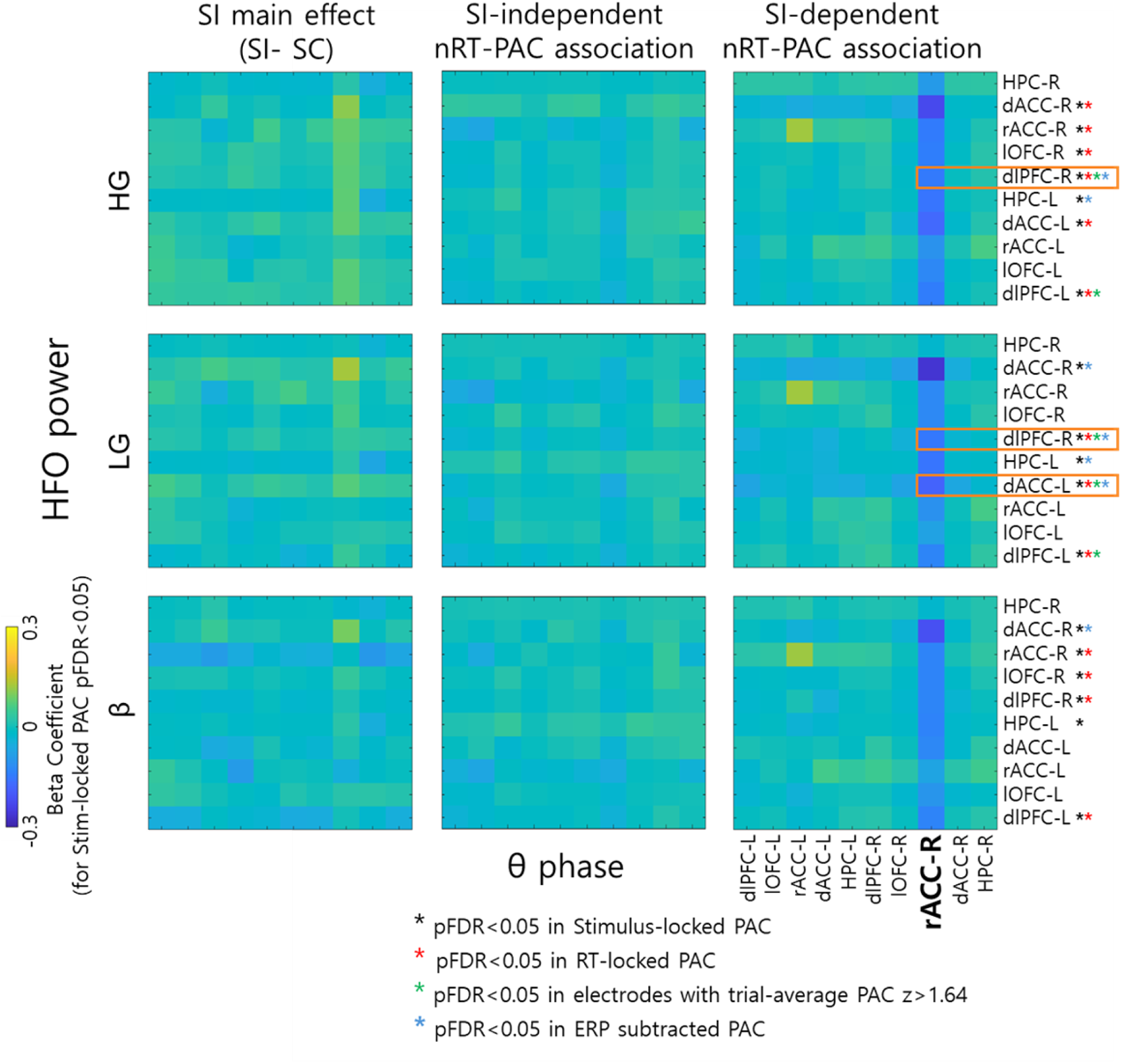
LMER testing the relationship between trial-by-trial PAC and nRT. We applied a LMER (*Methods*) to examine whether the relationship between trial-by-trial theta–HFO PAC for each regional pair (10 × 10 pairs) and nRT was modulated by CPE (negative interaction effect). Significant negative CPE × nRT interactions emerged only when the rACC-R served as the theta-phase source (pFDR < 0.05, corrected for 10 × 10 regional PAC pairs). There were no CPE-independent correlations between PAC and nRT, nor any PAC differences between the high- and low-CPE conditions. To further verify the robustness of the rACC-R PAC effects on control flexibility, we recomputed PAC using three complementary pipelines: RT-locked PAC, significant PAC (PACs above z > 1.64), and condition-specific ERP-subtracted PAC (red, green, and blue stars, respectively; black star for the original stimulus-locked PAC). theta–gamma PAC between rACC-R and dlPFC-R (theta–LG and theta–HG), as well as theta–LG PAC between rACC-R and dACC-L-two hubs of the conventional cognitive control network-consistently showed negative CPE interaction patterns across all four methods (orange box).

**Supplementary Figure 3.**
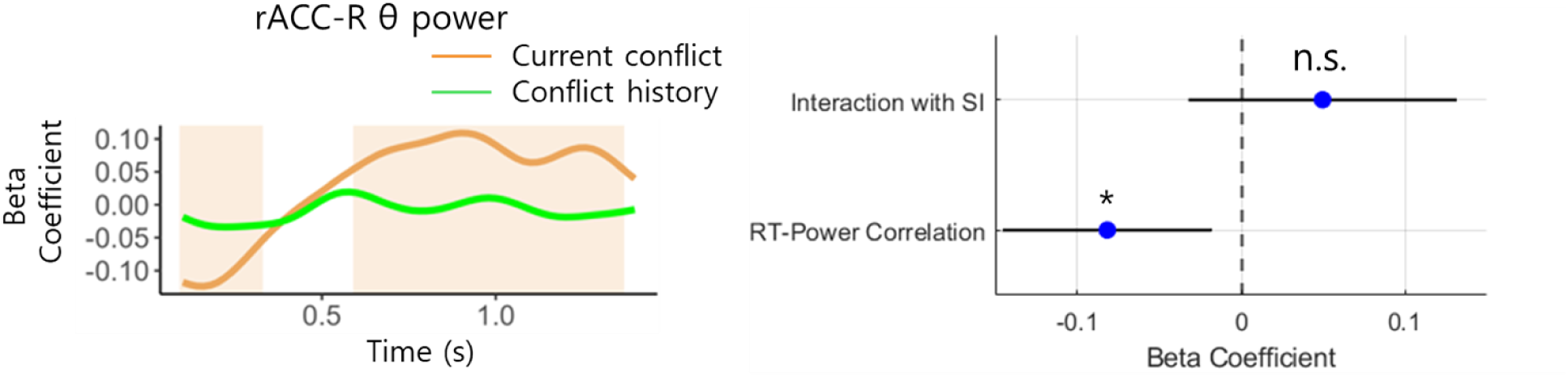
rACC-R theta power encoded current conflict, but not conflict history, in the cluster-based LMER analysis (left panel; *Supplementary Methods*). The subsequent theta Power–RT analysis (right panel; *Supplementary Methods*) on the current conflict–encoding cluster showed that rACC-R theta power reflected a general control mechanism, as stronger theta power was associated with faster nRT without a significant interaction effect in the LMER. * p<0.05, ** p<0.01, *** p<0.001.

**Supplementary Figure 4.**
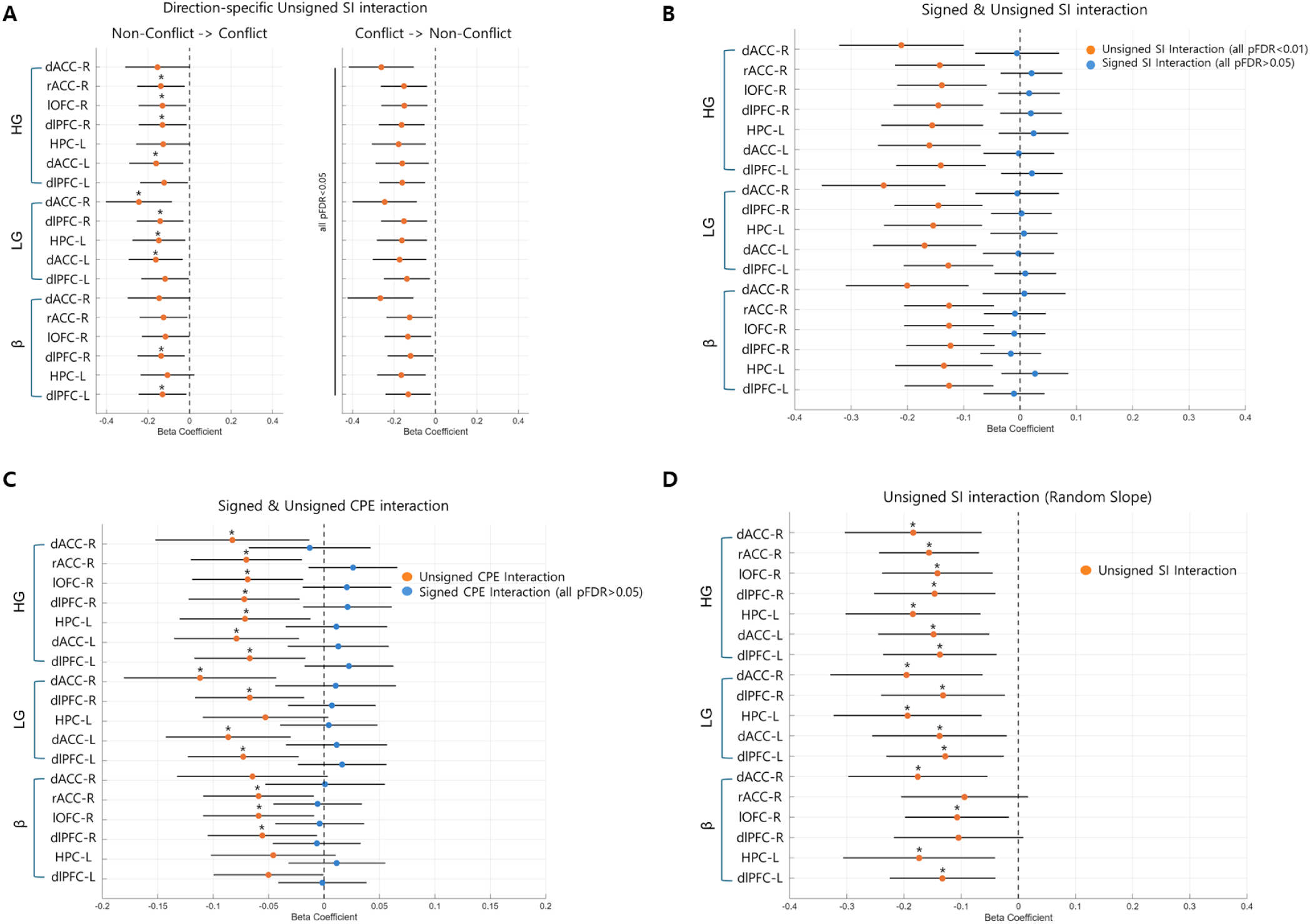
Supplementary validation analyses of the negative SI × nRT interaction effect on the RT–PAC relationship in rACC-R–centered PAC. Forest plots show LMER beta coefficients ± 95% confidence intervals for the interaction terms of interest across the 18 rACC-R–centered PAC region– frequency pairs from the primary PAC analysis (stimulus-locked PAC), grouped by amplitude band (HG, LG, and beta). The dashed vertical line indicates zero; asterisks denote FDR-corrected significance (*p*FDR < 0.05; corrected across the 18 selected region–frequency pairs from original PAC analysis). (A) direction-specific decomposition of SI showed negative interaction effects for both non-conflict → conflict and conflict → non-conflict transitions. (B) When unsigned and signed SI were modeled simultaneously, only unsigned SI × nRT remained significant. (C) When unsigned and signed CPE were modeled simultaneously, significant interaction effects were observed for unsigned CPE, but not signed CPE. (D) Random-slope models yielded a similar pattern, indicating that the main effect was robust to random-effects specification. Importantly, the three PAC pairs highlighted in the main manuscript—rACC-R theta–dlPFC-R LG PAC, rACC-R theta–dlPFC-R HG PAC, and rACC-R theta–dACC-L LG PAC—remained significant across all above analyses

**Supplementary Figure 5.**
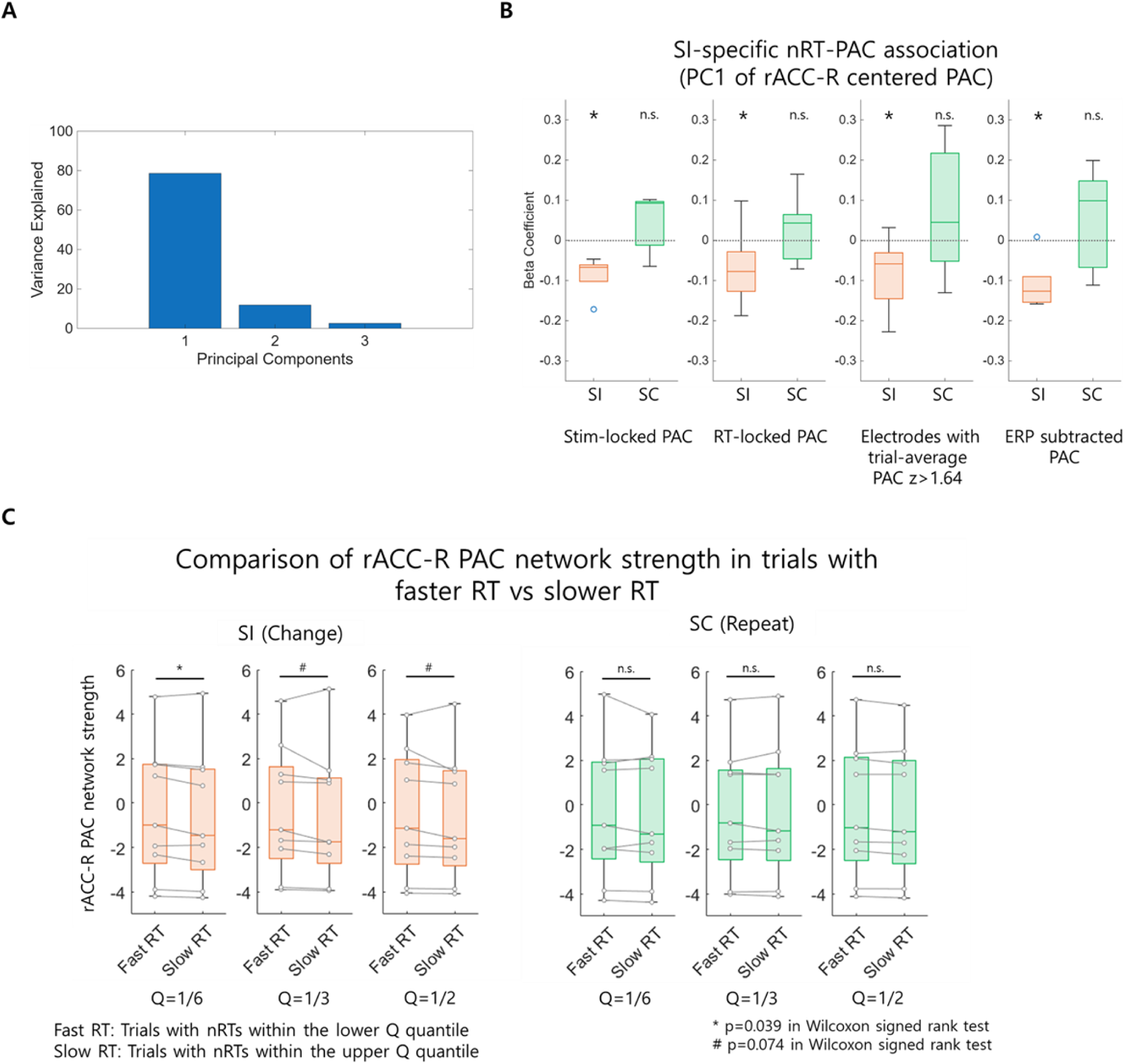
(A) To summarize the multi-regional PAC findings, we performed a PCA on the trial-wise rACC-R PAC matrix (# of trials × # of connected region–frequency pairs). The first principal component (PC1) typically explained 78% of the variance. (B) The PC1 loading, reflecting the overall strength of the rACC-R PAC network, exhibited the expected pattern of control flexibility: a negative nRT– PAC correlation under the SI condition (t = –2.53, coefficient = –0.10, pFDR < 0.05 in LMER, corrected for 4 methods) and no significant correlation under the SC condition (pFDR > 0.05) across all four methods. (C) To reduce noise inherent in trial-by-trial PAC estimates, we applied a quantile-averaging approach. Trials were binned into six quantiles based on nRT, and the mean rACC-R PAC network strength was compared between faster (upper sextile) and slower (lower sextile) trials under both SI and SC conditions. Average rACC-R PAC was higher in faster than slower trials under the SI condition (p = 0.039, Wilcoxon signed-rank test, n = 9), but not under the SC condition. Median- and tertile-based splits showed a similar, though weaker, trend (higher average PAC in faster vs. slower trials under the SI condition only, p = 0.074). * p<0.05, ** p<0.01, *** p<0.001, # p<0.1.

**Supplementary Figure 6.**
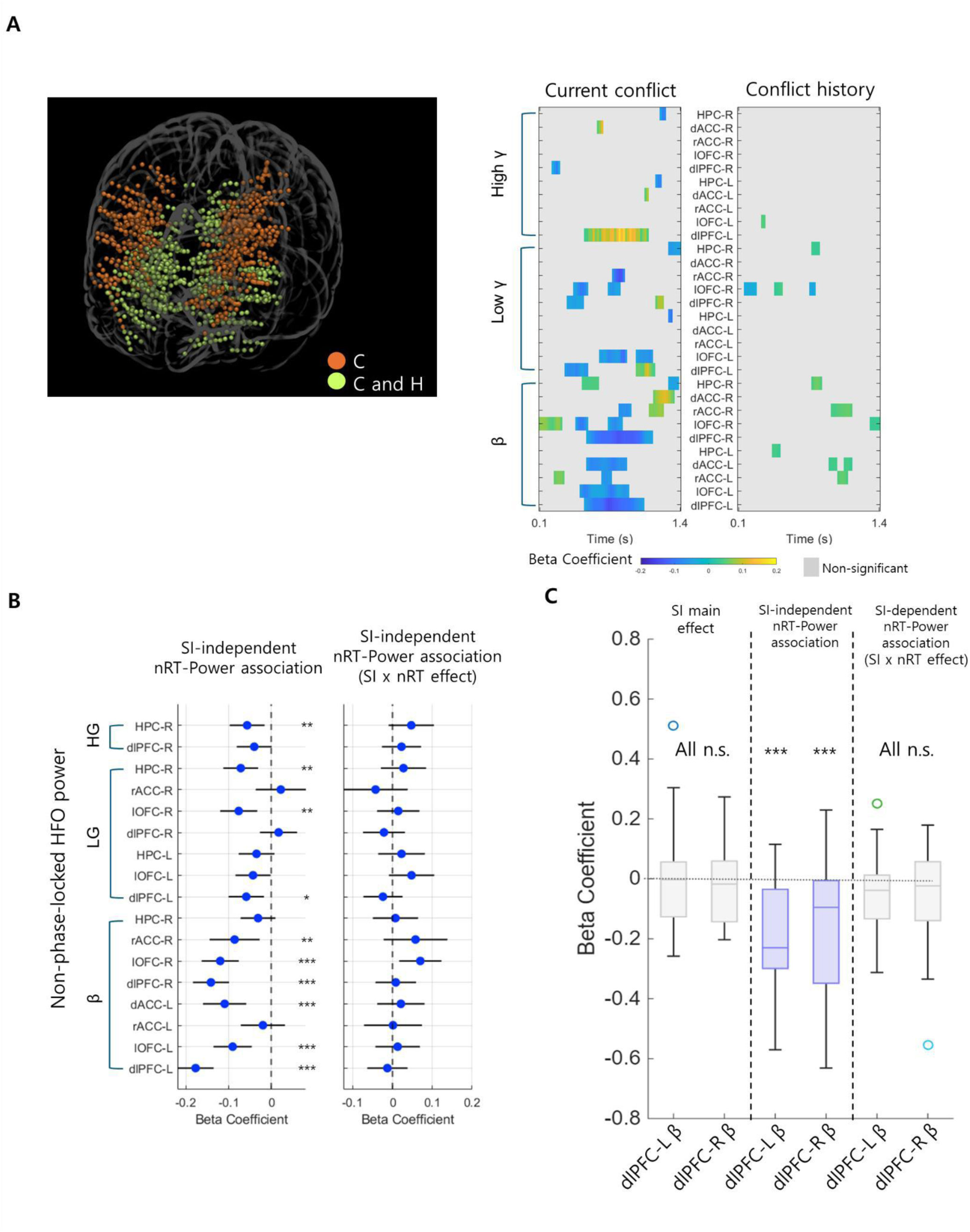
Local, non–phase-locked HFO power analysis. (A) Cluster-based permutation linear mixed-effects regression (cluster LMER) on local non–phase-locked HFO power identified significant time clusters (*p*FWE < 0.005) encoding current conflict only (orange) or both current conflict and conflict probability (CP; yellow), with CP reflecting recent conflict history. (C) vs. clusters encoding both current conflict and conflict history (C and H). (B–C) HFO Power–RT analysis (right panel; see *Supplementary Methods*) on the current conflict–encoding cluster revealed multiple CPE-independent RT–power correlations (pFDR<0.05; corrected for the number of significant clusters) without a significant CPE interaction, including dlPFC beta power. * p<0.05, ** p<0.01, *** p<0.001.

**Supplementary Figure 7.**
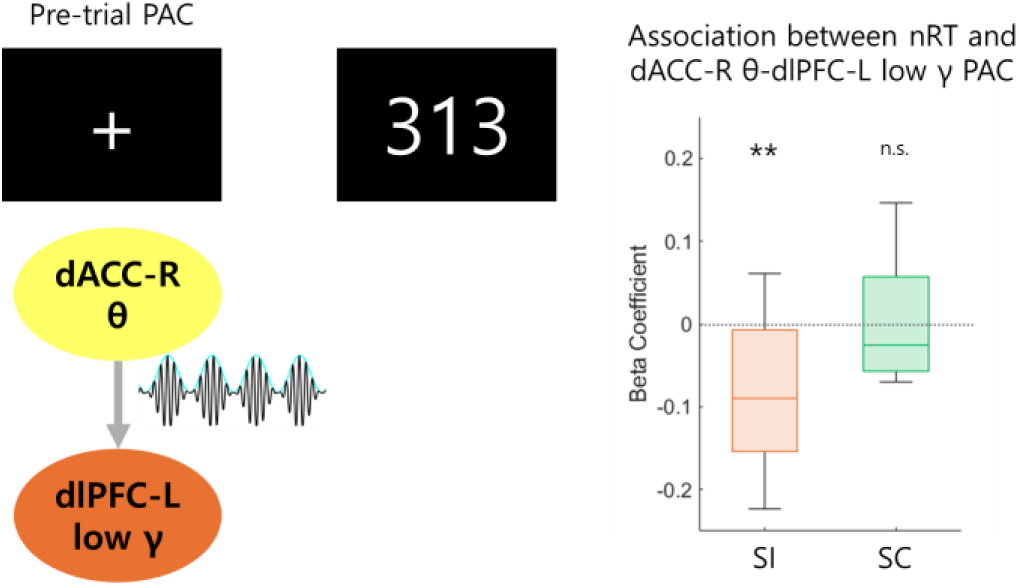
Effect of pre-trial dACC-R PAC on subsequent flexible adjustment. Pre-trial dACC-R–dlPFC-L theta–LG PAC was associated with faster nRT on the subsequent SI trial (coefficient = – 0.05, t = –2.86, pFDR = 0.023; FDR-corrected across 4 × 10 theta–low-gamma PAC pairs), but this association was absent when the upcoming trial was SC. * p<0.05, ** p<0.01, *** p<0.001.

**Supplementary Table 1.** Posterior-predictive model adequacy summary for the adaptive DDM. Observed behavioral summary statistics are compared with posterior-predictive simulations from the fitted adaptive DDM. Statistics include accuracy and upper/lower-bound RT distribution summaries; credible values indicate whether the observed statistic fell within the posterior-predictive distribution.

|  | stat | observed | mean | std | SEM | MSE | credible | quantile | mahalanob |
| --- | --- | --- | --- | --- | --- | --- | --- | --- | --- |
| 1 | accuracy | 0.98 | 1.00 | 0.01 | 0.00 | 0.00 | TRUE | 6.63 | 1.22 |
| 2 | mean_ub | 0.95 | 0.91 | 0.21 | 0.00 | 0.04 | TRUE | 64.56 | 0.16 |
| 3 | std_ub | 0.29 | 0.30 | 0.16 | 0.00 | 0.03 | TRUE | 61.60 | 0.04 |
| 4 | 10q_ub | 0.61 | 0.62 | 0.13 | 0.00 | 0.02 | TRUE | 58.39 | 0.06 |
| 5 | 30q_ub | 0.74 | 0.73 | 0.15 | 0.00 | 0.02 | TRUE | 62.63 | 0.06 |
| 6 | 50q_ub | 0.89 | 0.84 | 0.18 | 0.00 | 0.03 | TRUE | 69.52 | 0.26 |
| 7 | 70q_ub | 1.08 | 0.99 | 0.22 | 0.01 | 0.06 | TRUE | 74.72 | 0.43 |
| 8 | 90q_ub | 1.38 | 1.29 | 0.38 | 0.01 | 0.15 | TRUE | 68.49 | 0.25 |
| 9 | mean_lb | -1.00 | -1.23 | 0.64 | 0.05 | 0.47 | TRUE | 58.21 | 0.36 |
| 10 | std_lb | 0.35 | 0.36 | 0.39 | 0.00 | 0.16 | TRUE | 57.44 | 0.03 |
| 11 | 10q_lb | 0.63 | 0.90 | 0.53 | 0.07 | 0.36 | TRUE | 22.63 | 0.50 |
| 12 | 30q_lb | 0.73 | 1.01 | 0.56 | 0.08 | 0.40 | TRUE | 26.12 | 0.51 |
| 13 | 50q_lb | 0.89 | 1.15 | 0.62 | 0.07 | 0.46 | TRUE | 36.92 | 0.42 |
| 14 | 70q_lb | 1.25 | 1.34 | 0.74 | 0.01 | 0.55 | TRUE | 56.96 | 0.12 |
| 15 | 90q_lb | 1.53 | 1.64 | 0.97 | 0.01 | 0.96 | TRUE | 55.90 | 0.12 |

**Supplementary Table 2.** Intracranial electrode coverage by region in Dataset 1. Number of intracranial EEG electrodes and contributing participants are shown for each region of interest. Regions include bilateral dorsolateral prefrontal cortex, lateral orbitofrontal cortex, rostral ACC, dorsal ACC, and hippocampus.

|  | # of electrodes | # of participants |
| --- | --- | --- |
| **dIPFC_L** | 387 | 21 |
| **IOFC_L** | 116 | 18 |
| **rACC_L** | 35 | 12 |
| **dACC_L** | 75 | 17 |
| **HC_L** | 92 | 16 |
| **dIPFC_R** | 293 | 20 |
| **IOFC_R** | 133 | 20 |
| **rACC_R** | 13 | 9 |
| **dACC_R** | 41 | 14 |
| **HC_R** | 129 | 18 |
| **Total** | 1314 |  |

**Supplementary Table 3.** Tested VCVS Cog stimulation configurations. DBS OFF and all DBS ON research settings tested in the VCVS Cog parameter survey (Dataset 4) are shown. DBS ON settings used unilateral monopolar stimulation of left or right contacts 0–3 at 130 Hz, including 2 or 4 mA at 150 μs and 2 mA at 210 μs. DBS OFF sessions served as baseline comparison blocks; clinical-like settings selected for Dataset 4 analysis are summarized in Supplementary Table 4.

| Condition | Contact (0-3) | Amplitude (mA) | Pulse Width (ms) | Frequency (Hz) |
| --- | --- | --- | --- | --- |
| **DBS OFF** | N/A | N/A | N/A | N/A |
| **LEFT** | 0 | 2 mA | 150 ms | 130 Hz |
| **LEFT** | 0 | 4 mA | 150 ms | 130 Hz |
| **LEFT** | 1 | 2 mA | 150 ms | 130 Hz |
| **LEFT** | 1 | 4 mA | 150 ms | 130 Hz |
| **LEFT** | 2 | 2 mA | 150 ms | 130 Hz |
| **LEFT** | 2 | 4 mA | 150 ms | 130 Hz |
| **LEFT** | 3 | 2 mA | 150 ms | 130 Hz |
| **LEFT** | 3 | 4 mA | 150 ms | 130 Hz |
| **DBS OFF** | N/A | N/A | N/A | N/A |
| **RIGHT** | 0 | 2 mA | 150 ms | 130 Hz |
| **RIGHT** | 0 | 4 mA | 150 ms | 130 Hz |
| **RIGHT** | 1 | 2 mA | 150 ms | 130 Hz |
| **RIGHT** | 1 | 4 mA | 150 ms | 130 Hz |
| **RIGHT** | 2 | 2 mA | 150 ms | 130 Hz |
| **RIGHT** | 2 | 4 mA | 150 ms | 130 Hz |
| **RIGHT** | 3 | 2 mA | 150 ms | 130 Hz |
| **RIGHT** | 3 | 4 mA | 150 ms | 130 Hz |
| **RIGHT** | 0 | 2 mA | 210 ms | 130 Hz |
| **RIGHT** | 1 | 2 mA | 210 ms | 130 Hz |
| **RIGHT** | 2 | 2 mA | 210 ms | 130 Hz |
| **RIGHT** | 3 | 2 mA | 210 ms | 130 Hz |
| **LEFT** | 0 | 2 mA | 210 ms | 130 Hz |
| **LEFT** | 1 | 2 mA | 210 ms | 130 Hz |
| **LEFT** | 2 | 2 mA | 210 ms | 130 Hz |
| **LEFT** | 3 | 2 mA | 210 ms | 130 Hz |
| **DBS OFF** | N/A | N/A | N/A | N/A |

**Supplementary Table 4.** Clinical DBS settings and clinical-like VCVS Cog settings selected for Dataset 4. Chronic clinical DBS configurations are shown for each TR-OCD participant, together with the cathode contacts used to select clinical-like research stimulation settings from the VCVS Cog parameter survey. Because the research survey used monopolar stimulation settings, bipolar or multi-contact clinical configurations were approximated by selecting research settings involving the same cathode contact(s). For each selected contact, available research settings included 2 mA/150 μs, 4 mA/150 μs, and 2 mA/210 μs at 130 Hz. DBS OFF sessions served as the baseline comparison.

| Participant | Chronic clinical setting | Clinical cathode contacts used for matching | Clinical-like VCVS Cog settings selected | # ON sessions | Matching note |
| --- | --- | --- | --- | --- | --- |
| **VCVS001** | L: 2-/C+, 90 μs, 125 Hz, 7 mA;<br>R: 1-/C+, 120 μs, 125 Hz, 6 mA | L2, R1 | All L2/R1 monopolar research settings | 6 | Monopolar clinical settings; matched by cathode contact |
| **VCVS002** | L: 2,3-/C+, 210 μs, 130 Hz, 7 mA;<br>R: 2,3-/1+, 150 μs, 130 Hz, 7 mA | L2, L3, R2, R3 | All L2/L3/R2/R3 monopolar research settings | 12 | Dual cathodes; matched by cathode involvement |
| **VCVS003** | L: 0+ / 2,3-, 210 μs, 135 Hz, 6.1 mA;<br>R: C+ / 1,3-, 210 μs, 135 Hz, 10 mA | L2, L3, R1, R3 | All L2/L3/R1/R3 monopolar research settings | 12 | Multi-contact settings; matched by cathode involvement |
| **VCVS004** | L: 1-/C+, 180 μs, 125 Hz, 6 mA;<br>R: 1-/C+, 180 μs, 125 Hz, 6 mA | L1, R1 | All L1/R1 monopolar research settings | 6 | Matched by cathode contact |
| **VCVS005** | L: 3+ / 1,2-, 210 μs, 130 Hz, 10 mA;<br>R: 3- / 2+, 180 μs, 130 Hz, 10 mA | L1, L2, R3 | All L1/L2/R3 monopolar research settings | 9 | Bipolar/multi-contact settings; matched by cathode involvement |

## Supplementary Methods

### Time-Frequency Decomposition for the Local Power Analyses

Following our previous work (Dataset 1; Basu et al. ^28^), we analyzed neural activity in the 0.1 to 1.4 second window following image onset based on our previous work (Dataset 1 ^28^). This time window was chosen to capture the majority of stimulus-evoked processing and decision-making while excluding the earliest sensory transient. To examine whether the pre-stimulus period encoded conflict history information and whether DBS modulated this encoding, we also analyzed the −1.5 to −0.2 s window preceding image onset. This pre-stimulus window was chosen to assess history-related activity before onset of the next stimulus while avoiding contamination from immediate stimulus-locked activity. Time–frequency decomposition of the iEEG signal was performed using the FieldTrip toolbox ^72^, applying Morlet wavelet convolution to estimate spectral power across time and trials. Data were epoched from −2 to +3 s relative to stimulus onset, and wavelets were constructed with a width of 7 cycles per frequency to balance temporal and spectral resolution. Power was estimated from 2 to 150 Hz in logarithmic steps and sampled every 20 ms. For each electrode, power was averaged within canonical frequency bands: theta (4–8 Hz), beta (15–30 Hz), low-gamma (30–50 Hz), and high-gamma (70–150 Hz). Baseline correction was performed via z-scoring over the entire analysis window. Finally, power values were averaged across timepoints and frequencies within a band within 10 predefined regions of interest to extract trial-wise, region-level features for further analysis.

Notably, while theta-band power has been linked to cognitive control in prior studies ^28,29,73^, we did not include it in our local encoding analyses, as our primary interest was in phase dynamics—specifically, phase-amplitude coupling (PAC) between theta and HFO—rather than power-based theta modulation. However, we conducted theta power analyses for the rACC-R cluster as a control analysis to rule out the possibility that rACC-R theta power alone accounted for the observed relationship between rACC PAC and control flexibility. For DBS-related modulation of theta power in Dataset 1 and 3, see Basu et al. ^28^ and Widge et al. ^29^, respectively.

### Cluster-Based Permutation Tests

To investigate whether local power in high frequency bands encoded key cognitive control variables— namely, the conflict level on the current trial (termed “current conflict”) and conflict history (indexed as conflict probability or CP; computed from adaptive DDM model)—we performed a cluster-based permutation linear mixed-effects regression (cluster LMER) using the clusterperm.lmer package in R ^74^. To compute CP, we repeatedly sampled an adaptive-DDM parameter set for each participant from their posterior distribution and, for that set, simulated the trial-by-trial CP trajectory. We repeated this procedure 10,000 times and then averaged the resulting 10,000 simulated CP series to obtain the expected CP for every trial. Our linear mixed-effects model incorporated fixed effects for current conflict and conflict history, and blockNum as a covariate, and included random intercepts for each participant as below.

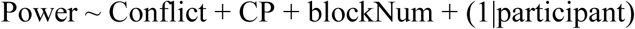

This analysis was applied to the power estimates at every time point across the 0.1–1.4 s window (sampled in 20 ms intervals). Family-wise error correction was achieved through permutation testing ^75^. Specifically, contiguous time points with p-values below a threshold of 0.05 were grouped into clusters, and the cluster-level statistics (e.g., the summed t-values) were compared against a null distribution generated from 1000 random permutations of the data. Clusters with a family-wise error corrected p-value (pFWE; corrected for every time point) below 0.005 were considered significant. Additionally, we tested whether pre-stimulus high-frequency power (from –1.5 to –0.2 s) encodes conflict history (CP) using the same cluster LMER approach. This approach enabled robust detection of time periods during which local high-frequency power significantly reflected either current conflict, conflict history, or both.

### Local Power–Behavior Analyses

To assess the behavioral relevance of these local power effects, we conducted trial-by-trial modeling on the significant clusters identified by the clusterLMER. We labeled the clusters for each region–frequency pair as either positive or negative. Clusters were labeled positive or negative according to the sign of the cluster-level LMER coefficient for the effect of interest. If multiple positive (or negative) time clusters were present within the same region–frequency pair, they were aggregated into a single positive (or negative) cluster. For each significant cluster, we extracted trial-wise HFO power and fit a linear mixed-effects model relating this power to RT and context. Specifically, we used a model of the form:

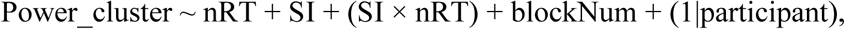

where *nRT* denotes the normalized log-transformed RT (z-scored within conflict vs. non-conflict conditions) and *SI* indicates sequential incongruency (high vs. low conflict-probability context). The model included the participant as a random intercept and block number as a covariate ^28^. This allowed us to test whether greater local high-frequency power predicted faster responses and whether any power–RT relationship was selectively stronger under high CPE (*SI*) conditions. Notably, while a generalized linear mixed model with a gamma distribution (to directly model RT’s skewed distribution and to maintain consistency with prior studies) was considered, we opted for the simpler LME with log-transformed RT due to the high computational cost of fitting a gamma GLMM across the large trial-by-trial PAC dataset and multiple region–frequency pairs. P-values were FDR-corrected for the number of regions in each frequency.

### Effects of ICS on Conflict History Encoding

Because an incorrect control prediction can interfere with the flexible encoding of new control information, we tested whether ICS improves CPE resolution by suppressing conflict-history signals under the high-CPE condition, when the predicted and actual control information mismatch. To examine this, we applied the following LMER model to the 10 clusters that encoded conflict-history information:

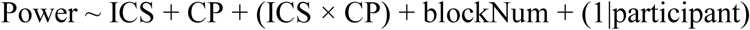

P-values were FDR-corrected for 10 clusters encoding conflict-history.

### Effects of ICS on General Control-related Local HFO Power

We tested whether ICS modulates HFO power associated with the general control signal (i.e., significant clusters showing nRT–Power correlations without CPE interaction; Supplementary Figure 6) using the following LMER model:

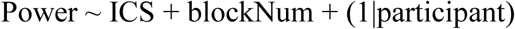

This analysis revealed enhanced bilateral dlPFC beta power (p_FDR<0.05, corrected for 10 clusters associated with the general control signal).

We then performed a mediation analysis to test whether the increased bilateral dlPFC beta power mediates the CPE-nonspecific decrease in RT—a hallmark of general control enhancement after ICS. Trial-by-trial right- and left-dlPFC beta power values were combined into a single bilateral dlPFC beta power index using PCA, with PC1 explaining 90% of the variance. Finally, we examined the correlation between enhancement of the general cognitive control parameter (ΔV_0_) and enhancement of bilateral dlPFC beta power.

## Supplementary Results

### Classical conflict-adaptation (Gratton) effect

Our primary behavioral analysis in the main manuscript used RT slowing following sequential incongruency (SI) as a marker of the adjustment burden imposed by a change in control demand. This differs from the classical conflict-adaptation (Gratton) framework ^32–34^, which focuses on the interaction between current-trial conflict and previous-trial conflict across the four trial-history combinations (CC, CI, IC, II). Because these two approaches are related but not identical, we quantified the classical Gratton effect in the non-psychiatric datasets to situate our SI-based framework relative to the more conventional behavioral literature.

Specifically, we fit trial-wise linear mixed-effects models predicting RT from current-trial conflict, previous-trial conflict, and their interaction. In these models, the critical Gratton term was the interaction between current-trial conflict and previous-trial conflict. A significant negative interaction indicates the classical pattern in which the effect of current conflict is attenuated following conflict trials.

In Dataset 1 (DBS OFF), the Gratton-style interaction was only marginal, although the effect was in the expected direction (coefficient = −0.02, *t*(5832) = −1.68, 95% CI [-0.04, 0.00], *p* = 0.094). In Dataset 2, the classical Gratton interaction was significant across all three task modalities (all *p* < 0.005).

Thus, the classical behavioral Gratton effect was robust in Dataset 2 but only marginal in Dataset 1. One likely reason for the weaker behavioral Gratton effect in Dataset 1 is task design ^28^. The MSIT in Dataset 1 was deliberately structured to minimize expectation effects associated with repetition of the same trial type, including a substantially lower probability of long same-type runs than in Dataset 2 MSIT (e.g., probability of four consecutive same-type trials: approximately 4% in Dataset 1 vs 13% in Dataset 2). This design feature would be expected to reduce the behavioral advantage typically observed when the same control state repeats across trials.

Importantly, our primary goal in the main manuscript was not to characterize classical direction-specific behavioral adaptation per se, but to identify the neural mechanism supporting efficient adjustment after control-state change. For that reason, we used the SI versus SC framework, which groups trials according to whether control demand changed or repeated, rather than distinguishing all four conflict-history conditions separately at the outset. We then directly tested whether collapsing across the two change directions was reasonable for the neural question and found that the negative nRT x SI interaction effect in rACC-centered PAC was present in both conflict → non-conflict and non-conflict → conflict transitions (*Supplementary Results*). Thus, even when the classical behavioral Gratton interaction was weak, the SI-related neural effect remained informative. We therefore interpret the rACC-centered PAC effect as reflecting a broader process of flexible adjustment to control-state change rather than as a conventional direction-specific conflict-adaptation effect.

### Identifying Locally Encoded Current Control vs History Information

Although we hypothesized that cross-frequency interactions—specifically, phase–amplitude coupling between the theta phase of ACC and high-frequency oscillations in LPFC —would support flexible control encoding ^15,17^, we also tested whether non–phase-locked high-frequency activity in these regions might locally encode control-relevant variables, including current conflict (Current Control information; Figure 1D) and recent conflict history (Conflict Probability, which influences control prediction; Figure 1D). These variables could potentially contribute to the computation of control prediction error (CPE; Figure 1D). To investigate this, we first examined high-frequency power (beta, LG, HG) in each region to determine whether any of them encoded these variables. This focus on high-frequency bands was further motivated by evidence linking these signals to decision-making processes: for example, beta-band power in LPFC has been associated with decision evidence ^76^, and gamma-band power in PFC has been tied to cognitive flexibility ^77^.Additionally, a recent study reported that local high-frequency power reflecting current conflict was associated with faster response times during cognitive control tasks ^31^. However, it remains unclear whether this signal directly supports cognitive flexibility (as in the pattern shown in Figure 2B) or reflects other types of control processes (e.g., general cognitive control; Figure 2C). To robustly detect any conflict- or history-related effects in local power, we performed a cluster-based permutation linear mixed-effects regression (“clusterLMER”; see Methods) ^74^ across the 0.1–1.4 s post-stimulus window (20ms time steps), with family-wise error correction across all time points (significance threshold *p*FWE<0.005).

From these clusterLMER analyses, we observed clusters where local high-frequency activity encoded current conflict, conflict history, or both. For example, bilateral dlPFC showed clusters encoding only current conflict across multiple bands (Figure 2C). In contrast, rACC, dACC, HC, and lOFC each contained clusters encoding both conflict history and current conflict (Figure 2C), suggesting that these areas might retain a memory trace of recent control demands and thus inform the internal estimate of conflict probability.

We next examined whether trial-by-trial high-frequency power in each identified cluster was correlated with RT and, critically, whether this relationship interacts with CPE as proposed in Figure 2B. Using a linear mixed-effects regression (see Methods), we tested whether the average beta/gamma power of each cluster is correlated with normalized log-RT (nRT; see Methods) on a trial-by-trial basis and if this power–nRT correlation was modulated by the CPE (high vs. low CPE, tested via SI x nRT interaction). The average power of many clusters encoding current conflict, especially in the beta band, were negatively correlated with nRT (all *p*FDR<0.05, corrected for number of regions; Figure 2C, right), indicating that stronger beta/gamma power predicted faster responses.

However—and crucially—no clusters exhibited a significant SI × nRT interaction in their local high-frequency power (all *p*FDR>0.05; Figure 2C, right). In other words, although local high-frequency activity encoding conflict correlated with faster information processing overall, in no region did power specifically track faster RTs in high-CPE (SI) trials. Thus, local high-frequency power is consistent with a general control mechanism in Figure 2B’s framework, rather than a flexible mismatch resolution process. Additionally, clusters encoding conflict history showed no correlation with nRT or the SI × nRT interaction.

This finding led us to hypothesize that inter-regional coordination, rather than any isolated local effect, might be key to resolving CPEs. Consequently, we turned to phase–amplitude coupling (PAC) analyses across sites, seeking a network-level mechanism for CPE resolution.

### Quantile-averaged approach to validate rACC-R PAC network

Additionally, we applied a quantile-averaged approach that could have lower noise than trial-by-trial PAC estimates (Supplementary Figure 3). Trials were binned into six quantiles based on nRT, and the mean rACC-R PAC network strength was compared between faster (upper sextile) and slower (lower sextile) trials in both SI and SC conditions. Average rACC-R PAC was higher in faster than in slower trials under the SI condition (p=0.039, Wilcoxon signed-rank test, n=9; Supplementary Figure 5C) but not under the SC condition. Median- and tertile-based splits showed a similar, though weaker, pattern (higher average PAC in faster vs. slower trials under the SI condition only, p=0.074; Supplementary Figure 5C).

### Effects of ICS on General Control-related Local HFO Power

Beyond flexibility, we probed ICS’ effects on general control that does not rely on CPE. ICS increased bilateral dlPFC beta power (all pFDR<0.05). Multilevel mediation showed that the bilateral dlPFC beta component (PC1) partially mediated the overall RT speeding under ICS (a×b=−0.02, p<0.001; Figure 4I). Across patients, ΔV_0_ (general control enhancement after ICS) correlated with dlPFC beta enhancement (Spearman rho=0.89; p<0.05; N=6; Figure 4G), suggesting that augmenting dlPFC beta (e.g., via IC WM tracts connecting to dlPFC) can support CPE-independent control, even though ΔV_0_ increased in only 4/6 patients.

### Additional validation analyses of the main SI × nRT effect

#### Supplementary validation of the main SI × nRT effect 1: Direction-specific decomposition of sequential incongruency

To test whether the main rACC-centered PAC effect depended on a specific direction of control-state change, we decomposed sequentially incongruent (SI) trials into two transition types: non-conflict → conflict (C→I) and conflict → non-conflict (I→C). We then re-estimated the LMER models underlying the main SI-related negative interaction effect on the RT–PAC relationship separately for these two transition directions within the 18 rACC-R–centered PAC region–frequency pairs identified in the primary analysis, using the same model specification as in the original analysis:

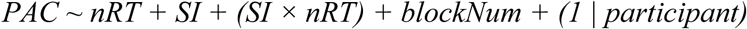

This analysis was motivated by the possibility that SI might reflect qualitatively distinct processes depending on whether control demand increased or decreased. If the rACC-R–centered PAC effect were specific to deploying control in response to increased conflict, it should be present primarily in C→I transitions. Conversely, if it reflected disengagement from conflict, it should be present primarily in I→C transitions.

Across the 18 primary rACC-centered PAC pairs, negative SI-related interaction effects on the RT–PAC relationship were observed in both C→I and I→C transitions, with coefficients in the same direction and of broadly similar magnitude (Supplementary Figure 4A). Ten of the 18 region–frequency pairs showed significant negative interaction effects in **both** transition directions (all *p*FDR < 0.05), including the three PACs emphasized most strongly in the main manuscript—rACC-R theta–dlPFC-R LG PAC, rACC-R theta– dlPFC-R HG PAC, and rACC-R theta–dACC-L LG PAC. In addition, all 18 pairs showed significant negative interaction effects in the C→I direction, whereas the remaining 8 I→C effects were generally still negative but did not all reach corrected significance. Thus, the neural interaction of interest was not restricted to only one direction of control-state change. Instead, it was consistent with a broader role for rACC-centered PAC in reconfiguring control state when recent demand changed.

#### Supplementary validation of the main SI × nRT effect 2: Direction-specific decomposition of sequential incongruency Simultaneous signed and unsigned sequential-incongruency regressors

To test whether the main PAC effect—namely, the SI-related interaction effect on the RT–PAC relationship centered on rACC-R PAC—was better explained by non-directional, unsigned control-demand change than by direction-specific control change, we re-estimated the PAC models including both unsigned SI and signed SI regressors simultaneously. This analysis was performed for the 18 region–frequency pairs with rACC-R as the theta phase-providing region that showed significant negative unsigned-SI interaction effects in the original stimulus-locked PAC analysis reported in the main manuscript. The following LMER was used:

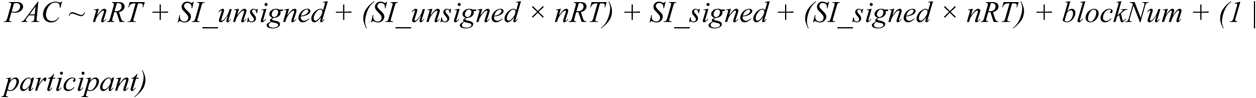

Unsigned SI coded whether control demand changed relative to the previous trial irrespective of direction, whereas signed SI coded the direction of that change (non-conflict → conflict trials were coded as 1, conflict → non-conflict trials were coded as −1, and no-change trials were coded as 0).

This analysis directly addressed whether the rACC-centered PAC effect was more consistent with flexible adjustment of control driven by an unsigned mismatch / surprise signal or by a direction-specific control-deployment signal. If the latter were true, the key interaction should be better captured by signed SI. If the former were true, the key interaction should remain associated with unsigned SI even after accounting for the signed regressor.

Consistent with our hypothesis, the critical interaction effect on the RT–PAC relationship remained associated only with the unsigned SI regressor (all *p*FDR < 0.05 across the 18 region–frequency pairs; Supplementary Figure 4B), whereas the corresponding interaction with signed SI was not significant (all *p*FDR > 0.05). Thus, the rACC-R PAC–RT relationship was modulated by the presence of a control-state change rather than by its direction. This supports the interpretation that the network dynamics of interest are more closely related to non-directional mismatch / surprise than to a direction-specific control-deployment account.

#### Supplementary validation of the main SI × nRT effect 3: Continuous signed and unsigned CPE regressors from the adaptive DDM

SI/SC is a task-level categorical contrast indexing whether control demand changed relative to the previous trial. In the main manuscript, we showed that RT slowed when control demand changed (SI vs SC), and that this task-level effect could be computationally explained by greater unsigned control prediction error (CPE), which in turn requires greater adjustment.

Several theoretical frameworks propose that the ACC detects valence-neutral surprise or mismatch signals in the form of control prediction error, thereby encoding not only whether a mismatch occurred, but also its magnitude. If the ACC—particularly the right rostral ACC, which showed a significant SI-related interaction effect in the main analysis—tracks control-demand mismatch in the form of unsigned CPE and supports flexible adjustment when that mismatch is large, then we would expect a corresponding interaction effect of CPE on the RT–PAC relationship, analogous to the interaction effect observed for SI.

To test this hypothesis, we extracted trial-by-trial unsigned CPE estimates from the adaptive DDM for each participant and used these quantities as regressors in the LMER models. To test the alternative possibility that the flexible-control effect was better explained by signed CPE rather than unsigned CPE, we also included signed CPE in the same model:

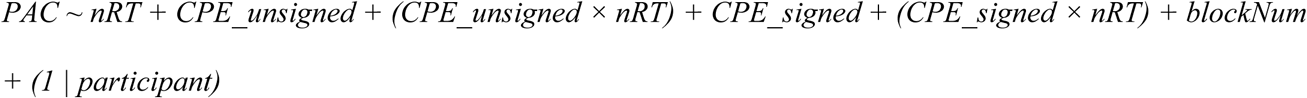

Unsigned CPE indexed the magnitude of discrepancy between predicted and observed control demand, whereas signed CPE preserved the direction of that discrepancy. This analysis therefore tested whether the main rACC-R PAC effect generalized from the observable task-level manipulation to the model-derived latent variables, and whether it was better captured by unsigned rather than signed mismatch. This analysis was performed for the 18 region–frequency pairs with rACC-R as the theta phase-providing region that showed significant negative unsigned-SI interaction effects in the original stimulus-locked PAC analysis reported in the main manuscript.

Consistent with our hypothesis, the interaction effect on the RT–PAC relationship was associated only with the unsigned CPE regressor (*p*FDR < 0.05 in 14 of the 18 region–frequency pairs, including rACC-R theta– dlPFC-R LG/HG PAC and rACC-R theta–dACC-L LG PAC; Supplementary Figure 4C), whereas the corresponding interaction with signed CPE was not significant (all *p*FDR > 0.05). Thus, the rACC-R PAC– RT relationship was modulated by the magnitude of control-demand mismatch rather than by its direction. This pattern is consistent with prior ACC frameworks, including the PRO model, in which valence-neutral surprise signals track discrepancies between expected and observed events and support adaptive adjustment.

#### Supplementary validation of the main SI × nRT effect 4: Random-slope sensitivity analyses

Following our prior work ^28,29^, the primary LMER analyses in the main manuscript were fit with random intercepts by participant. To test whether the principal findings depended on this random-effects specification, we performed sensitivity analyses using by-participant random slopes for the principal predictors, where estimable. Specifically, we fit the following model, where SI denotes unsigned sequential incongruency:

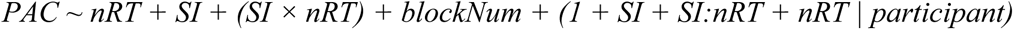

For the set of LMERs corresponding to the principal rACC-R–centered findings (18 rACC-R theta phase– amplitude coupling region–frequency pairs), we asked two questions: (1) whether including random slopes materially changed the main SI-related interaction effects on the RT–PAC relationship, and (2) whether the added complexity of the random-slope models was justified by improved model fit relative to the random-intercept-only models.

Including random slopes yielded a broadly similar pattern of results. Specifically, significant negative SI interaction effects on the RT–PAC relationship remained in 14 of the original 18 region–frequency pairs, including the key rACC-R theta–dlPFC-R LG/HG PAC pairs and the rACC-R theta–dACC-L LG PAC pair (*p*FDR < 0.05 in 14/18 pairs; Supplementary Figure 4D). Moreover, the simpler random-intercept models provided better fit than models including both random intercepts and random slopes (ΔBIC = −56.32 for no-slope minus slope models, 95% CI [−60.70, −51.94], *t*(17) = −27.15, *p* < 0.001). Thus, the use of random-intercept models in the primary analyses reflected a preferable bias–variance tradeoff rather than an arbitrary analytic choice.

### Relationship of the flexibility parameter ω to global RT

Because the flexibility parameter ω scales the effect of control-demand mismatch on drift rate, we examined whether it might also reflect more general differences in response speed. This issue is particularly relevant because internal capsule stimulation (ICS) produced a main effect on RT.

In our adaptive DDM, the parameters most directly intended to capture general response speed are baseline drift rate and decision threshold, whereas ω scales how strongly control-demand mismatch influences drift rate. As a supportive sanity check, we therefore tested whether participant-level mean log RT was explained by fitted drift rate, threshold, and ω. Mean log RT was associated with drift rate and threshold, but not with ω (*p* = 0.8389), consistent with the view that ω does not primarily reflect baseline response speed.

We next asked whether the stimulation-related change in ω could be better explained as a more general, context-independent change in response speed. In Dataset 1, we explicitly compared models in which ICS affected: (1) general control only, via baseline drift rate; (2) general control plus a CPE-independent increase in ω; and (3) general control plus a CPE-dependent increase in ω. If the stimulation effect were explained primarily by a more general, context-independent change in response speed captured through ω, stronger support would be expected for the model with CPE-independent increases in ω. Instead, the best-fitting model was the one with CPE-dependent increases in ω together with a separable general-control component, indicating that the ICS effect was better explained as mismatch-sensitive flexibility enhancement rather than as a purely global RT effect.

Consistent with this dissociation, in Dataset 3 the flexibility-related parameter (Δω), rather than the general-control parameter (ΔV₀), was associated with clinical response. Together, these results support interpreting ω primarily as a parameter indexing the mismatch-sensitive component of control rather than general response speed.

### Sensitivity analysis addressing temporal spread of the low-theta Morlet wavelet

To assess whether the main stimulus-locked PAC effects might have been driven by pre-trial contributions from the low-frequency Morlet wavelet tail, we repeated the stimulus-locked PAC analysis using a later post-stimulus window (0.6–1.6 s instead of 0.1–1.4 s). This later window was chosen because, at 4 Hz with 7 cycles, the Gaussian width parameter is s=n/(2πf)≈0.279s, such that 0.6 s ensures that the 2SD tail does not contain timepoints before stimulus onset. We then recomputed the SI × nRT interaction for the 18 rACC-R theta-centered PAC pairs identified in the original stimulus-locked analysis. Results remained highly similar: 16 of the 18 PAC pairs remained significant after FDR correction, including the 3 key PACs involving rACC-R theta phase with dlPFC-R LG, dlPFC-R HG, and dACC-L LG. These findings indicate that the main stimulus-locked PAC effects were robust to substantial delay of the analysis window and were therefore unlikely to be driven primarily by pre-trial activity.

